# Genomic Evidence for Males of Exceptional Reproductive Output (ERO) in Apes and Humans

**DOI:** 10.1101/2025.06.20.660753

**Authors:** Xiaopei Wang, Hongpu Chen, Lingjie Zhang, Mei Hou, Yang Gao, Xuemei Lu, Pengfei Fan, Miles E. Tracy, Liying Huang, Haijun Wen, Yongsen Ruan, Shuhua Xu, Chung-I Wu

## Abstract

It is widely suspected that a small percentage of males have exceptional reproductive output (ERO) but progeny numbers of males are rarely measurable, even in humans. If we define the variance ratio of reproductive-output in males and females as *α’* = *V*_*M*_/*V*_*F*_, the ERO hypothesis would predict *α’* >>1. Since autosomal, X, and Y chromosomes are found in males 1/2, 1/3, and 100% of the time, their DNA diversities can inform about *α’*. For example, autosomal and Y-linked diversities are governed, respectively, by (*V*_*M*_+*V*_*F*_)/2 and *V*_*M*_. When comparing the chromosomal diversities, *α’* appears to be near 20 for chimpanzees and orangutans, and 1-10 for gorillas. The exception is bonobos with *α’* < 1. In humans, the extensive genomic data are coupled to a theory, developed herein, that can filter out selection influences on Y-linked diversities. Hence, the estimation of *α’* is rigorous, yielding values near or above 20, depending on the population. When *α’* >10, the presence of ERO males is very likely. These analyses can be applied more generally to species with XY sex determination.

## Introduction

In sexually reproducing species, the two sexes may evolve divergent reproductive strategies toward breeding success [1-3], defined as each individual’s progeny *(K)* reaching maturity. Obviously, the mean breeding success for males and females, *E*_*M*_*(K)* and *E*_*F*_*(K)*, should be the same. In contrast, the variance of male breeding success, *V*_*M*_*(K)*, is likely larger than that of females, *V*_*F*_*(K)*.

An extreme form of male reproductive strategy may be the phenomenon of ERO males. It has often been speculated that a small percentage of males in many species have an outsize contribution to the gene pool [4-6]. However, the difficulties in measuring male breeding success have hampered studies of social-sexual behaviors and, more generally, of sexual selection. Here, we are interested in great apes including humans. Despite extensive field studies on primates [7-12] as well as anthropology, sociology, and behavioral biology on humans [3, 13], the breeding successes of individual males are usually uncertain.

If we define *α’* = *V*_*M*_*(K)*/*V*_*F*_*(K)*, field studies of monkeys have suggested *α’* to be between 1.5 and 6.5 in*Macaca* or *Mandrillus* species [14, 15]. Of greater challenges are ERO males whose reproductive successes (i.e., their K values) are outliers in the biological sense. (In statistical terms, outliers are males from the tail end of a highly kurtotic K distribution.) For example, if 1% of males are outliers with *K* = 40 while keeping *E(K)* ∼ 1 (per parent), *V*_*M*_*(K)* and *α’* would increase by > 15-fold. In other words, *K* should be highly kurtotic if *α’* is large, say > 10 (see Discussion for further details). ERO males are therefore few and likely to be missed in field studies. They are occasionally known, or suspected, in human societies [5, 16-18] but can be further investigated by estimating *α’* more broadly across species with XY sex determination.

This ratio *α’* can be obtained from the polymorphism data of the Y, X, and autosomes (A) as they respectively spend 100%, 1/3, and 1/2 of the evolutionary time in males. Note that the ratio of mutation rate between sexes, *α* (without the prime), has been estimated using the same logic [19]. The difference is that the mutation rate is estimated from the level of species divergence whereas *α’* must be estimated from the within-species polymorphism.

The goal of this study is to obtain the levels of neutral polymorphisms among A, X, and Y to estimate *α’*. However, the relative levels of X/A and Y/A are simultaneously affected by selection and sex-dependent drift. While studies focus on either selection [20-22] or sex-dependent drift [4, 23, 24], it is clear that both are operative. In particular, selection is stronger on X- and Y-linked than on autosomal genes because recessive mutations are exposed to selection and X-Y recombination is absent. This exposure can reduce the X/A and Y/A polymorphism ratios [22, 25, 26].

Since our goal is to estimate *α’* from DNA sequence data, it would be best to filter out *cleanly* the effect of selection on DNA diversities. As each Y chromosome is a non-recombining haplotype, Y’s are amenable to the estimation of the true neutral diversity whereas the task is more difficult for X. For this reason, the Y/A ratio, after the correction for different selective pressures, should be more reliable for estimating *α’* than X/A (see Results). When properly done, the estimation of *α’* may provide new perspectives into the social-sexual behaviors of the great apes. Nevertheless, cautions must be taken when applying the approach developed here to other taxa with chromosomal sex determination due to the many nuances that could make the estimation challenging. Such nuances include the different selective pressures on sex chromosomes vs autosomes [22, 25, 26] as well as the history of X vs Y evolution [27, 28]. Finally, because the dynamics of the XY and ZW systems mirror each other, one may wish to extend the analyses to the ZW systems. In Discussion, we suggest the refinements that will be needed for such extensions.

## Results

In PART I, the theory for estimating *α’* = *V*_*M*_*(K)*/*V*_*F*_*(K)* based on the data of neutral nucleotide diversity is provided. In the analyses of non-human apes in PART II, we use the measure of polymorphism that is relatively insensitive to selection, *θ*_*w*_ [29], when estimating the *diversity ratios* between X, Y and autosomes. In contrast, another commonly used statistic, *θ*_*π*_, is strongly influenced by selection [22, 30] in this application. Nevertheless, there may still be residual influences of selection in the *θ*_*w*_ data. Because the X vs. Y-linked reductions have opposite effects on *α’*, an MSE estimation is presented to balance the biases. In PART III, the extensive human data motivate the development of *θ*_*0*_ estimation that is the theoretical limit of selection-free diversity. The caveat is that the procedure is applicable only to the non-recombining Y-linked mutations. Generally speaking, the estimations of *α’* offer a clear picture of the evolution of the social-sexual behaviors in apes and humans.

### PART I General theory for estimating the *V*_*M*_*(K)*/*V*_*F*_*(K)* ratio

Let *K* be the progeny number of each individual. The mean values of *K* for males and females are generally the same, or *E*_*M*_*(K)*=*E*_*F*_*(K)*. The variances of *K* are *V*_*M*_*(K)* and *V*_*F*_*(K)* and are generally believed to be *V*_*M*_*(K) > V*_*F*_*(K)*[11, 31]. The ratio *α’* is our main interest:

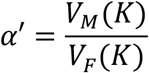

This ratio can be estimated from the polymorphisms of Y, X, and autosomal (A) sequences since they are in males 100%, 1/3, and 1/2 of the time. The level of DNA polymorphism is governed by the parameter *θ* (=4*N*_*e*_*μ*) where *N*_*e*_ is the effective population size and *μ* is the mutation rate. *N*_*e*_ and *μ* are both different for all three chromosomes. For convenience, we denote the mutation rate of X, and Y chromosomes as xμ, and yμ, relative to the autosomal mutation rate μ. *N*_*e*_ is a function of the actual population size, *N*. The exact function would depend on the biology and evolution of the population. In the context of sexual reproduction, we use *N*_*e*_ = *N/V(K)*, which was first obtained by [32, 33] and further elaborated in [34, 35]. For the three sets of chromosomes, under standard assumptions of diploidy and 1:1 sex ratio, *N*_*e*_*(Y)=N(Y)/V*_*Y*_*(K), N*_*e*_*(X)=N(X)/V*_*X*_*(K)* and *N*_*e*_*(A)=N(A)/V*_*A*_*(K)* where *N(A):N(X):N(Y)* = 4:3:1 and *V*_*[Y,X or A]*_*(K)*’s are as follows:

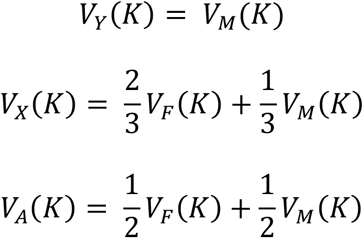

From the observed polymorphisms, *θ* (=4*Nμ/V(K)*) can be estimated. The Supplementary Note 1 shows that *α’* can be obtained 3 ways:

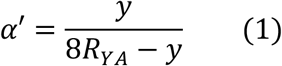

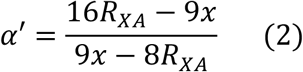

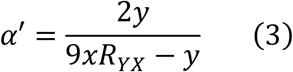

where *R*_*YA*_, *R*_*XA*_, and *R*_*YX*_ are, respectively, 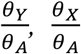 and 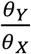. The ratios can be estimated by the relative level of polymorphisms among the three chromosomes.

Two main challenges have to be met in applying Eqs. (1-3) on genomic data to obtain *α’*. The first one is the extreme sensitivity of the *α’* estimates to measurement errors in mutation rate and polymorphism. Fig. 1A shows the dependence of *α’* on *R*_*YA*_ and *y*. Note that the parameter spaces for *α’* < 1 and *α’* reaching infinity are much larger than the space for *α’* in the feasible range (> 1 but < infinity). The yellow band for the interval of [10, 20] is narrow, as shown, while the space for [20, infinity] is too small to show (see Fig. 1A legend). Hence, if the biology is such that *α’* > 10, the data would likely show *α’* < 10 or approaching infinity. Fig. 1B shows the same sensitivity in the X vs. A comparison. The second challenge is natural selection (both positive and negative) which would reduce the level of polymorphism [36-38]. Interestingly, under-estimating polymorphism on X and Y would have opposite effects on the estimated *α’* values (Fig. 1A vs. 1B).

**Figure 1.**
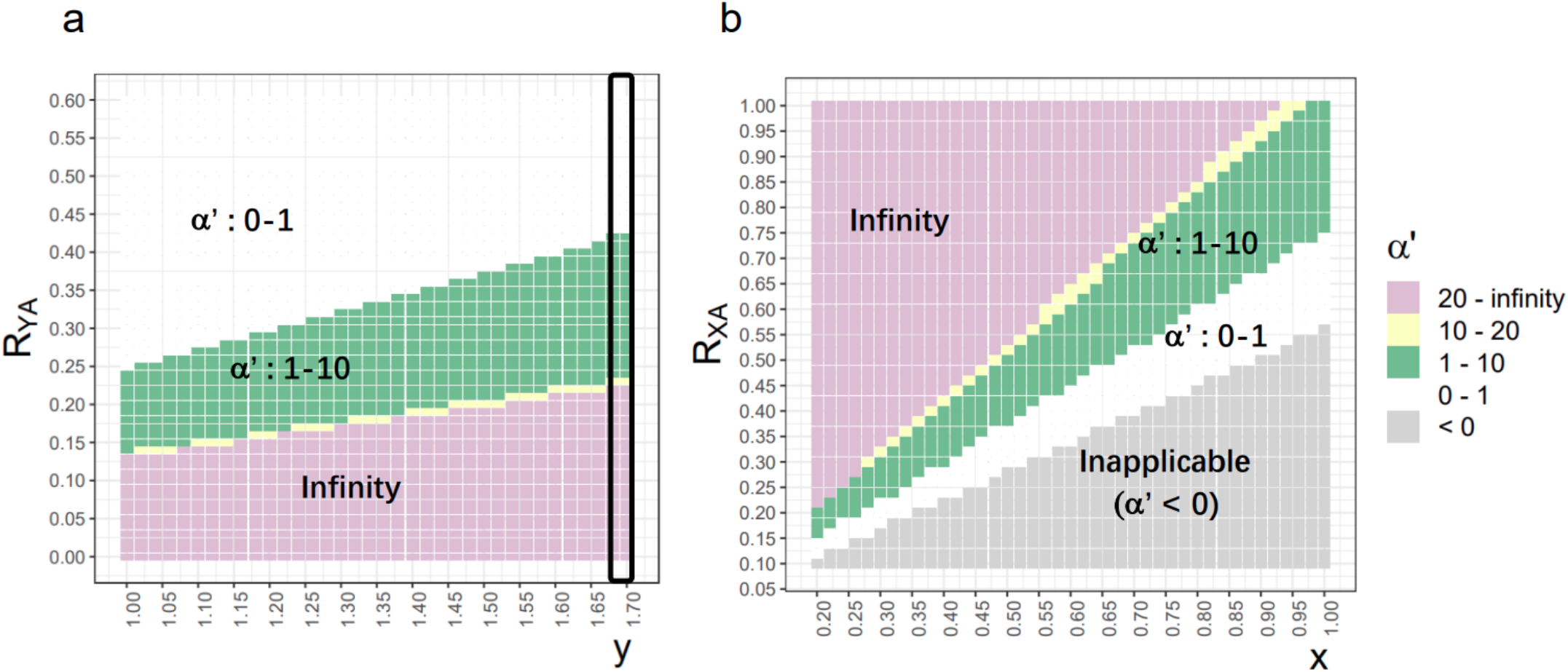
The Dependence of *α’* on the relative diversity (*R*) and mutation rate (*x, y*) - The *α’* values are shown by different colors in the heatmap. (a) Estimation based on the Y-A comparison. The X axis is the *y* value, which is the relative mutation rate of Y and autosomes. The Y axis is *R*_*YA*_ (= *θ*_*Y*_/*θ*_*A*_). Note that the yellow band for *α’* in the interval (10, 20) is much smaller than the green band for (1, 10). The band for *α’* in the interval (20, infinity) is even thinner than the yellow band and is hence merged with the pink zone of infinity. The black rectangle represents the distribution of *α’* at *y* = 1.68. Note that *α’* increases as *R*_*YA*_ decreases. (b) Estimation based on the X-A comparison. The X axis is *x* value and the Y axis is *R*_*XA*_. Contrary to (a), *α’* decreases as *R*_*XA*_ decreases. In both (a) and (b), the yellow strip shows the narrow parameter space for *α’* between 10 and infinity.

To meet the challenges, we use the statistics that mitigate the influence of selection and measurement errors in PART II. In PART III, an explicit model that incorporates selection and chromosome-dependent *N*_*e*_ is developed for human datasets.

### PART II. Genomic polymorphisms in non-human apes

To estimate *α’* by Eqs. (1-3), we need to obtain the relative levels of polymorphism, *R*_*YA*_, *R*_*XA*_, and *R*_*YX*_, as well as the relative levels of mutation rate, *x* and *y*. Since these quantities are based on neutral variants, noncoding region sequences are used. The values of *x* and *y* in humans and chimpanzees have been reported as *x* = 0.77 - 0.80 and *y* = 1.60 - 1.68 [39, 40]. A salient property of population genetics is that the neutral rate of divergence between species can be formulated as *2Nμ(1/2N)* = *μ* [33, 41]. The rate is hence a function of the mutation rate only (*x* and *y* here) and does not depend on *N* or *N*_*e*_. We shall use the values most commonly used long-term species divergence estimate [39] of *x* = 0.77, *y* = 1.68. It should be noted that most of the published numbers are within a tight range [40, 42]. Furthermore, the paired (x, y) estimates of different studies vary in ways that affect the estimation of *α’* minimally (see Fig. 1).

The extraction of *θ’*s from the sequence data demands close attention. As stated, the *θ* value of each chromosome has to represent its neutral level. However, the observed diversities on Y, X, and A may not reflect the neutral levels equally. In particular, both X- and Y-linked diversities are expected to be lower than autosomes due to the stronger influence of selection on two fronts: 1) the hemizygosity in XY males exposes recessive mutations; and 2) the hemizygosity also reduces recombination, thus facilitating hitchhiking with other mutations on the same X or Y chromosome. Between X and Y, genes on Y are fully linked but there are far fewer fitness-related mutations. This disparity arises from the Y chromosome’s lower gene density and their preservation may enhance male fitness but with limited overall fitness impact [26, 43, 44].

Among the many statistics for *θ* [29, 36, 37, 45, 46], those giving more weight to low-frequency variants are less sensitive to selection (see Eq. (4) below), including *θ*_*w*_ [29] and *θ*_*1*_. While *θ*_*w*_ is more commonly used, *θ*_*1*_ (only singletons are considered) is much closer to the neutral state than other variants [37]. The most commonly used statistics, *θ*_*π*_ [22, 30, 47], would be dictated by the mid-range frequencies (near 0.5) where selection influence would predominate.

As noted in Fig. 1A-1B, the estimated *α’* values are very sensitive to even slight variations in the observed diversities (*R*_*YA*_, *R*_*XA*_, and *R*_*YX*_). To alleviate the sensitivity, we combine the three estimates of Eqs. (1 - 3) by identifying *α’* that yields the minimum MSE (Mean Squared Error; see Eq. (5) in Methods). Note that a decrease in Y-linked polymorphism would lead to over-estimating *α’* but the opposite trend is true for X-linked polymorphism. Therefore, the minimum MSE estimates of *α’* should buffer against under-estimation of sex-linked (both X and Y) polymorphisms.

The data of non-human great apes were compiled by Hallast, et al. [30] from various publications [25, 48] on the 4 species - bonobo, chimpanzee, gorilla, and orangutan (Supplementary Materials and Table S1). We use *θ*_*w*_ to estimate the minimum MSE estimation of *α’* in Table 1. The basic information is shown in the first 5 columns. In the three *θ*_*w*_ columns, X-linked diversity is usually lower than the autosomes and, crucially, Y-linked diversity is always lower than autosomes and by a larger margin.

**Table 1.** *α’* estimates for non-human great apes (*θ*_*w*_)

| Non-human great Apes |  |  |  |  |  |  |  |  |  |  |
| --- | --- | --- | --- | --- | --- | --- | --- | --- | --- | --- |
| | n<br>(Y, X, A) | $\theta_w(Y)$<br>($\times 10^{-3}$) | $\theta_w(X)$<br>($\times 10^{-3}$) | $\theta_w(A)$<br>($\times 10^{-3}$) | $Z(YX)$ | $Z(XA)$ | $Z(YA)$ | MSE<br>$\alpha'$ | Expanded<br>n(Y, X, A) | MSE<br>$\alpha'$ |
| Range for Z<br>with $\alpha' > 1^*$ | - | - | - | - | < 0.33 | > 0.75 | < 0.25 | - | - | - |
| Bonobo | (4, 4, 26) | 0.492 | 0.446 | 0.867 | 0.506<sup>†</sup> | 0.669 | 0.338 | 0.570 | (7<sup>‡</sup>, 24<sup>§</sup>, 26) | 0.574 |
| Chimpanzee |  |  |  |  |  |  |  |  |  |  |
| Western | (9, 7, 10) | 0.034 | 0.533 | 1.624 | 0.029 | 0.426 | 0.012 | 4.98 | (77<sup>‡</sup>, 7, 10) | 4.140 |
| Eastern | (3, 3, 12) | 0.1442 | 1.044 | 1.446 | 0.063 | 0.938 | 0.059 | 25.9 | (6<sup>‡</sup>, 10<sup>§</sup>, 12) | 3.762 |
| Nigeria-Cameroon | (4, 4, 20) | 0.213 | 0.8181 | 0.703 | 0.119 | 1.511 | 0.180 | Inf | - | - |
| Gorilla |  |  |  |  |  |  |  |  |  |  |
| W lowland | (7, 7, 54) | 0.1376 | 0.7219 | 1.555 | 0.087 | 0.603 | 0.053 | 4.53 | (7, 50<sup>§</sup>, 54) | 5.366 |
| E lowland | (3, 3, 18) | 0.0653 | 0.0921 | 0.714 | 0.325 | 0.168 | 0.054 | 1.11 | - | - |
| Mountain | (3, 3, 14) | 0.0007 | 0.2211 | 0.652 | 0.001 | 0.440 | 0.001 | 6.07 | - | - |
| Orangutan |  |  |  |  |  |  |  |  |  |  |
| Sumatran | (4, 4, 10) | 0.0221 | 1.2282 | 2.0089 | 0.008 | 0.794 | 0.007 | 23.49 | (4, 9<sup>§</sup>, 10) | 22.356 |
| Bornean | (2, 2, 10) | 0.1354 | 0.5569 | 1.4369 | 0.111 | 0.503 | 0.056 | 3.35 | (2, 9<sup>§</sup>, 10) | 3.294 |

The X and Y data are from [30]; the autosomes data from [25], except the Gorilla autosome data are from [48]. The sample sizes for each chromosome are denoted as n(Y, X, A).
\* This row gives the expected range of Z values for $\alpha' > 1$ . The expected value is based solely on the nominal numbers of A, X and Y in the population, given that $Z(YX)$ , $Z(XA)$ , and $Z(YA)$ are the ratios of the normalized nucleotide diversity with chromosome-specific mutation rate.
<sup>†</sup> Red letters indicate cases of Z values yielding $\alpha' < 1$ .
As explained in the main text, $Z(YX)$ , $Z(XA)$ , and $Z(YA)$ are biased in different ways. If we examine each column separately, $Z(YX)$ and $Z(YA)$ columns are significantly different from the predicted values under $\alpha' =$ 1. Whereas $Z(XA)$ data by themselves do not reject the null hypothesis of $\alpha' = 1$ .
Expanded sample sizes are provided in the 10<sup>th</sup> column, with new data noted as follows: <sup>‡</sup>data from [85];
<sup>§</sup>data from [25].

These patterns are more intuitively readable in the next three *Z* columns. The *Z* ratios, *Z(YA), Z(XA)*, and *Z(YA)*, are represented by 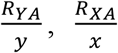, and 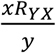, respectively. Hence, the *Z* values would reflect the chromosome polymorphism, normalized with chromosome-specific mutation rate, relative to copy number. For example, *Z(YA)* < 0.25 would mean *α’* > 1, since the copy number of Y is 1/4 of that of autosomes. Those entries that do not show *α’* > 1 are shown in red letters. One can see that *Z(XA)* values are often shown in red. As stated, both X- and Y-linked polymorphisms are underestimated since natural selection is more effective on hemizygous chromosomes. The MSE estimates of *α’* in the third column from right would mitigate these biases as under-estimation of X- and Y- diversity affects *α’* in the opposite directions.

Because the data come from multiple sources, we choose a subset that is most consistent in quality (such as read length, coverage and clarity in annotation). As a result, the sample size is sometimes on the lower side. In the last two columns, the stringency of data filtering is relaxed to allow for a larger sample size (Table 1). The alternative *α’* estimates are largely concordant with those of the stringent set; however, one glaring discrepancy exists. The augmented *α’* for Eastern chimpanzee is reduced from 25.9 to 3.76. There are both biological and technical reasons for such a large difference. Nevertheless, both estimates still show *α’* > 1.

Overall, the bonobo is exceptional as all *α’* estimates are < 1. The species has a matriarch social-sexual society [8, 10, 49]. It would seem plausible that *V*_*F*_*(K)* would be unusually large and *V*_*M*_*(K)* would be relatively small. All other *α’* estimates are > 1. In gorillas, *α’* estimates are smaller than those in chimpanzees. Perhaps, with the harem system [9, 50-52], gorillas may not have as many outlier males of high reproductive output as chimpanzees may have. Nevertheless, we should caution again that *α’* estimates are very sensitive to measurement errors.

Fig 1A-1B shows that the parameter space for *α’* in [1, 10] or > 20 is large. In contrast, the parameter space for [10 - 20] is only a tiny strip shown by the yellow band in Fig. 1. Indeed, among chimpanzees, gorillas and orangutans, 5 *α’* values fall in the range of [1-10] and 3 fall above *α’* > 20. If we interpret *α’* as the degree of highly biased reproductive success, the result may suggest a bimodal distribution of male reproductive behavior among these species, or even among populations. However, this would not be the correct biological interpretation. If there is no measurement errors, *α’* values could be quite common in the interval of [10, 20]. In fact, α’ >> 20 would seem biologically implausible for many species.

### PART III. Analyses of human polymorphisms incorporating selection and drift

To reduce any potential batch effects in analyzing multiple data sets from different resources, we obtained whole-genome deep sequencing data of Africans, Europeans, and East Asians from the Human Genetic Diversity Panel (HGDP) [53]. For the A, X, and Y diversities, we randomly selected 84 males of each population to obtain the polymorphism data. Only one haploid of the autosomes is used from each male, thus guaranteeing an identical sample size of *n* = 84 for each chromosome. The same procedure is then applied to the data analysis of the 1000 Genomes Project [54] data with *n* = 46. Details can be found in Supplementary Materials.

We will show mathematically that the low-frequency portion of the mutation spectrum is closest to the expected pattern under no selection. Therefore, both *θ*_*w*_ [29] and *θ*_*1*_ [37] are used. While both positive selection and negative selection can influence *θ* estimation, PART III will focus on negative selection. Positive selection is of lesser concern and can be more easily interpreted after all results have been presented (see Supplementary Note 5).

#### 1. Estimation of α’ based on θ_w_ and θ_1_

In Table 2, we present the analysis of human polymorphisms based on the HGDP sample of *n* = 84. (Results from a second dataset of KGP with n = 46, shown in the last column, will be discussed later.) The upper part of Table 2 is based on *θ*_*w*_ as in Table 1. Here, the *α’* values in the three populations are close to those of gorillas but lower than those of chimpanzees. The lower part of Table 2 presents the analysis of *θ*_*1*._ In comparison, *θ*_*1*_ exhibits higher values, indicating a higher degree of correction for the selection effect. Consequently, *Z*(*XA*) is larger by *θ*_*1*_ than by *θ*_*w*_, leading to a larger *α’*. With *θ*_*1*_, the lower part of Table 2 indeed reveals *α’* > 1 by all three measures, *Z(XA), Z(YA)* and *Z(YX)* and the MSE estimates of *α’* in humans fall in the range of 10 – 20.

**Table 2.**
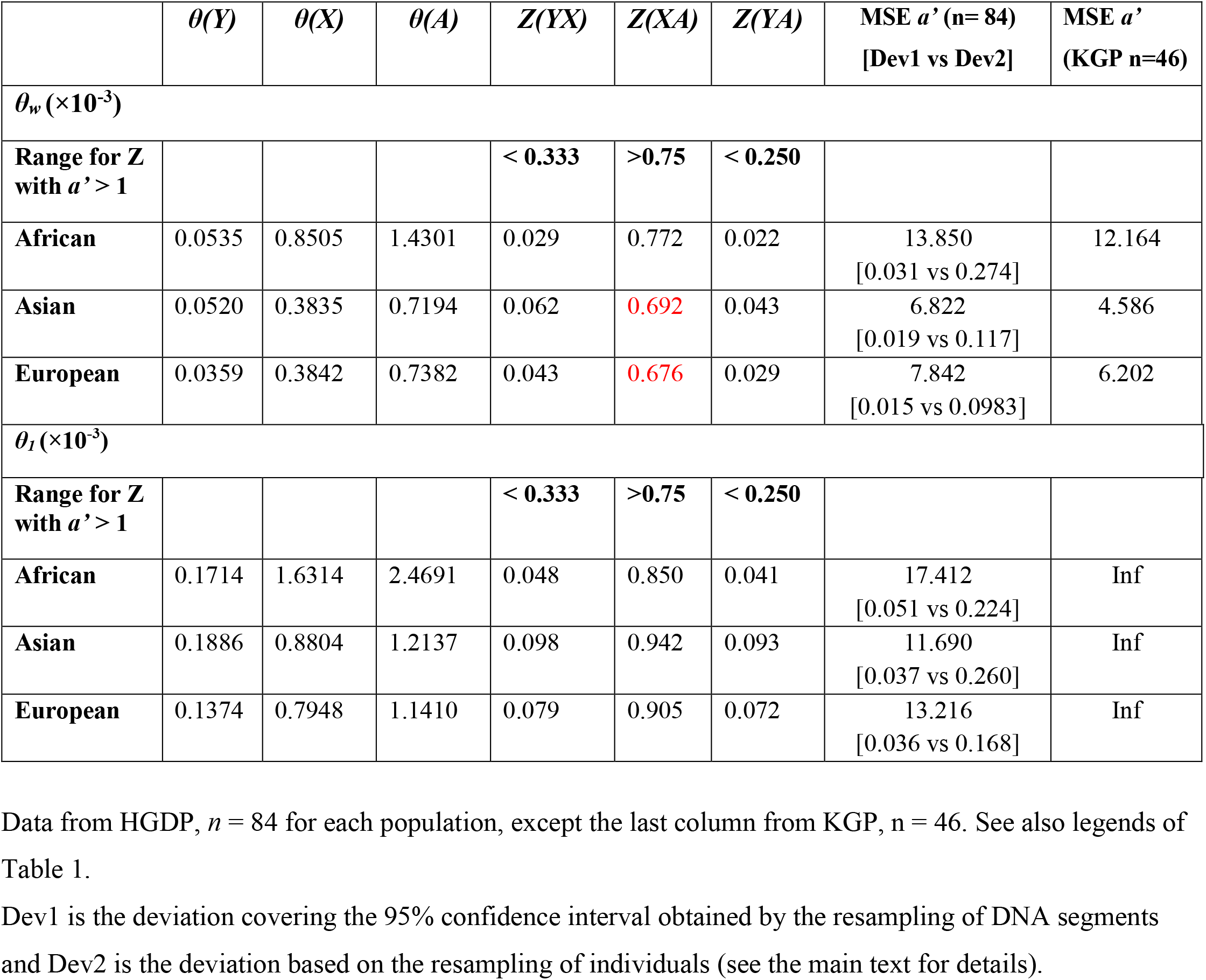
*α’* estimate in humans based on *θ*_*w*_ and *θ*_*1*_.

For the confidence intervals, we perform two different bootstrapping procedures on the larger dataset of n = 84). For this purpose, we may resample either the DNA segments or the individuals, yielding deviations as dev1 and dev2, respectively. While the commonly used procedure is to resample the same number of times from the original sample with replacement [55, 56], the two procedures are very different in scale.

There are more than 3 million 1Kb segments to be resampled while there are only 84 individuals for bootstrapping. Details of the bootstrapping procedure are given in Methods.

As shown in the second to the last column of Table 2, Dev1 (from resampling DNA segments) is very small, most likely due to the sheer number of DNA segments. Dev2 (from resampling individuals) is substantially larger with 84 individuals to be resampled. Nevertheless, the confidence intervals are small by either scheme. Importantly, resampling individuals introduces substantial bias due to the nature of the *θ* statistics. In particular, *θ*_*1*_ relies on variants that occur in only one individual. In resampling, such singleton variants can be easily missed or appear more than once, and thus not counted. For that reason, we report Dev2 values as confidence intervals surrounding the estimates of the original estimates.

Regardless of how bootstrapping is carried out, it is still based on the information embedded in the original sample. For that reason, we use a separate but smaller dataset (KGP) to calculate independent estimates for *α’*. As shown in the last column of Table 2, these independent estimates are fairly close to the HGDP-based estimations. Considering that these two datasets are both based on samples from multiple populations within Africa, Asia and Europe, we can be confident that the results of Table 2 are robust.

#### 2. The extrapolation of variant frequency to 0

It would be desirable to cleanly filter out the influence of negative selection on Y-linked polymorphisms. If the variant frequency approaches 0, then Y/A should ideally represent the neutral ratio. In PART III, we let *x* be the variant frequency of both Y- and A-variants. This subsection presents only Eq. (4) of the theory for *θ*_*0*_ where *x* approaches 0. This would be a check of how closely *θ*_*1*_ represents the neutral value. Details are shown in Methods.

Let Φ′(*x*) be the site frequency spectrum of Y-linked variants that are under selection of intensity *s* and Φ(*x*) be that of the autosomal variants without selection. The selective coefficient, *s* of SFS represents the joint effect of selection on all mutants on the Y (could be as low as one single mutant), averaged over time. The ratio, F(*x*) = Φ^*^(*x*)/Φ(*x*), is obtainable from the polymorphism data. We assume *N* = 7500 for human autosomes [57, 58]. For the Y chromosome, *N’* = *ηN* where *η* = 0.5 in the ideal population. Hence, *η* < 0.5 is a measure of *V*_*M*_*(K)*. Also, we use the relative mutation rate ratio of Y-to-A = 1.68 [39]. When *x* is in the low frequency range (< 0.1), the derivation in Methods show:

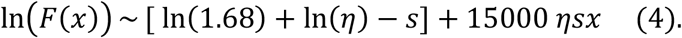

Importantly, Eq. (4) is a linear function of *x* with a slope of 15000*ηs*. It can be seen from Eq. (4) that the low-frequency portion of the spectrum (*x* ∼ 0) is minimally affected by selection when the 15000*ηx* approaches 0.

The goal is to obtain F(*x* ∼ 0) from Eq. (4), which will be equated with *R*_*YA*_ in the calculation of *α’* by Eq. (1). Since ln[F(*x*)] is linear with *x*, we should be able to obtain F(*x* ∼ 0) from the observed data [i.e., F(*i/n*)] if we know the slope. (In practice, we only do the extrapolation from F(1/*n*) as its level of polymorphism is highest and sufficiently reliable.) To obtain the slope, we calculate one slope for each pair of ln[F(*i/n*)] and ln[F(*j/n*)] for 0 ≤ *i* < *j, j* ≤ 4. We also use a dummy F^*^(*x* ∼ 0) = *ηy =*0.5 *y*, which is the theoretical maximum for F(*x* ∼ 0). For each dataset, there should be 10 slopes for five data points 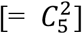. Each slope is used to calculate a different F(*x* ∼ 0) estimate, thus yielding 10 different *α’* estimates from one dataset (see methods).

#### 3. Estimation of α’ based on θ (x ∼ 0)

For each human population, there are 18 estimates of *α’*, 9 from each dataset. Because the estimation is sensitive to measurement errors, we present each *α’* estimate that represents one set of parameter measurements (i.e., the slope of Eq. (4)). The spread of these *α’* estimates should more faithfully show where the actual *α’* falls. The details are presented in Table S4.

The overall results are summarized in Fig. 2 which includes those of Tables 1 and 2. In Fig. 2, great apes in general show *α’* > 1 based on *θ*_*w*_. The parameter space for *α’* falling in (1 - 10), (10 - 20) and (> 20) are displayed according to the size of the segment. (Note that the expanded dataset, collected from various sources, yield concordant results but with one excetion; see legends of Fig. 2). The bonobo is a singular exception with both *α’* < 1. Gorillas have all four estimates in the interval of (1 - 10). For chimpanzees and orangutans, the *α’* estimates fall on the two sides of the (10 - 20) interval, suggesting that the true values may indeed be within, or slightly to the right of (10 - 20).

**Figure 2.**
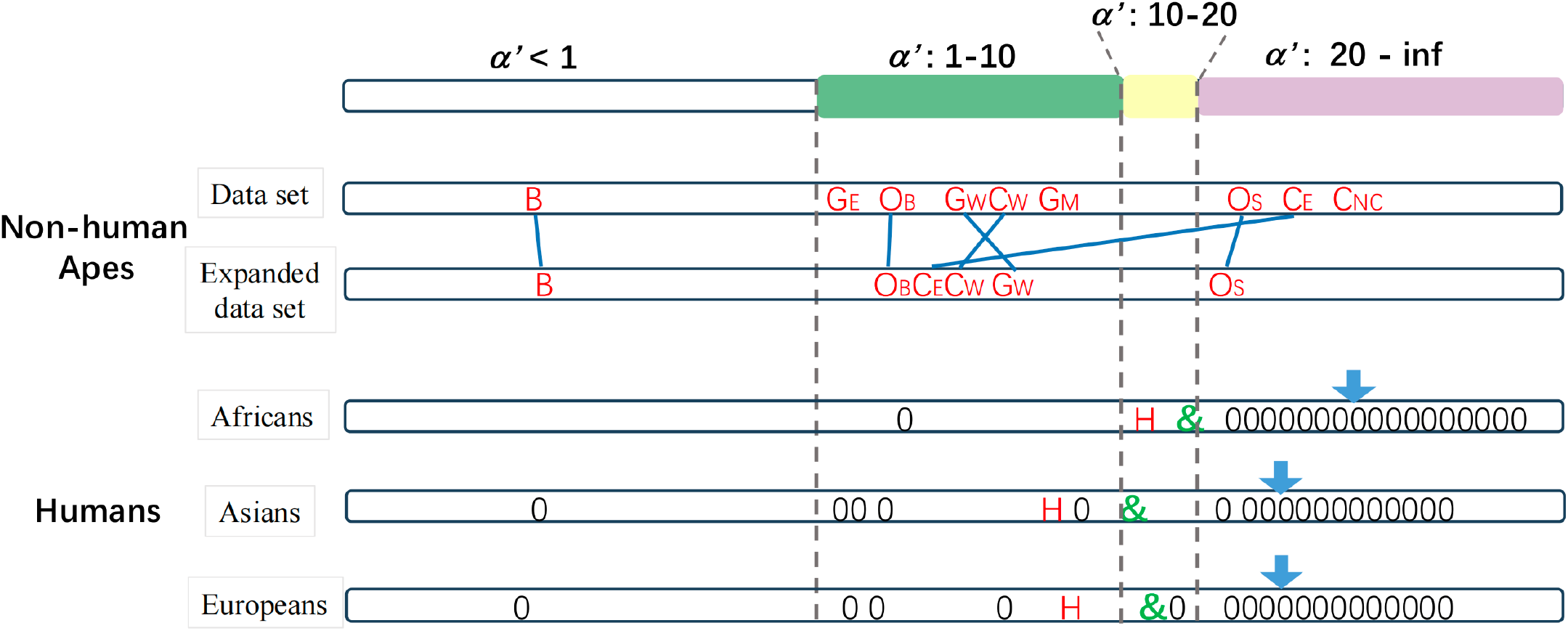
Summary of the estimation of *α’* = (*V*_*M*_*(K)/V*_*F*_*(K)*). The estimation of *α’* has been done by procedures based on various *θ* statistics, each of which is accompanied by bootstrapped confidence intervals. Here, we further present all *α’* estimates obtained from different datasets and procedures in one single figure to portray the broader range of *α’* estimation. The estimates of *α’* fall into 4 intervals (< 1, 1 - 10, 10 - 20 and > 20). The estimates based on *θ*_*w*_ are shown by red letters: Bonobo (B), Chimpanzee: Western (C_W_), Eastern (C_E_), Nigeria-Cameroon (C_NC_); Gorilla: Western lowland (G_W_), Eastern lowland (G_E_), Mountain (G_M_); Orangutan: Sumatran (O_S_), Bornean (O_B_) and Human. For non-human great apes, the *α’* estimates by expanded sample size corresponding to the last column of Table 1, are shown as the expanded data set. Two *α’* estimates for the same species are connected by grey lines. For humans, the green-letter &’s are based on *θ*_*1*_. The bootstrapping values of H’s and &’s are provided in Table 2 and Table S3, respectively, along with estimates from another dataset (see the text for further detials.) The black-letter 0’s are *θ*_*0*_ estimates based on Eq. (4). Each 0 represents an estimate based on a different subset of the observed diversity values. The estimates are placed, roughly in order, only in intervals up to 20. When an *α’* estimate falls above 20, there is little resolution. Hence, 0’s in that interval should all be read simply as *α’* > 20.

For humans, *α’* estimates are shown by *θ*_*w*_ (red letter H in Fig. 2), *θ*_*1*_ (green letter &) and *θ*_*0*_ (0’s in Fig. 2; based on Eq. (4)). First, the red-lettered *α’* estimates based on *θ*_*w*_ are lower than the green letters based on *θ*_*1*_. Since the other apes are analyzed by *θ*_*w*_, their *α’* values could be underestimated as well. Second, *α’* estimates by *θ*_*0*_ (with the arrowhead showing the median of 0’s) are moderately higher than by *θ*_*1*_. Third, *α’* estimates for Africans are consistently higher than for Asians and Europeans. The conservative conclusion is that, with the exception of the bonobos, great apes including humans have *α’* near, or larger than, 10. Given the small parameter space shown in Fig. 1A, the resolution is reliable up to *α’* ∼ 10, whereas there is little resolution between *α’* ∼ 20 and infinity (see Fig. 1 legend).

Despite the difficulties in obtaining the reliable confidence intervals for *α’*, the statistical patterns of Fig. 2 are robust. The H and & symbols are supported by bootstrapping on the larger data set (second to the last column, Table 2). Furthermore, a smaller independent dataset yields concordant estimates (last column, Table 2). Finally, these conservative estimates, as expected, fall in the tail end of a cloud of estimates based on *θ*_*0*_ (the 0’s). In Discussion, we will connect the high *α’* estimates to the “ERO males” hypothesis.

## Discussion

The conclusion of ERO males, based on the genomic data, can now be treated as a hypothesis. Importantly, the basis of comparisons should be the “adjusted” level of polymorphism after the different strengths of selection among X, Y and autosomes have been accounted for. Among the measures, *θ*_*v*,_ *θ*_*w*,_ *θ*_*1*_ and *θ*_*0*_, the last one (i.e., *θ*_*0*_ estimate of Y-linked polymorphism) is most appropriate. Its feasibility is due to the complete linkage that enables the estimation of selection on the entire Y. (In contrast, comparisons involving X are less accurate because its *θ*_*0*_ estimate cannot be obtained due to recombination).

Given that the equations for the XY and ZW systems are mirror images, the approach seems easily transferrable to the ZW system whereby females are heterogametic, as in birds and butterflies. However, in such systems, the confounding effects of selection would pose far more difficulties. In particular, most published diversities in ZW species are based on *θ*_*v*_’s[59, 60], the least desirable measures. It seems obvious that further developments of theories as done in PART III for Y will be necessary for the Z and W chromosomes. We shall now discuss three types of tests: i) field observations, ii) social-sexual conditions, and iii) other possible tests.

1. Field observations - The overall conclusions are broadly consistent with the general understanding of the social-sexual behaviors of great apes [3, 9]. Bonobo is the only species with *α’* < 1 as calculated by *θ*_*w*_. Since the estimation based on *θ*_*w*_ would yield a lower *α’* than *θ*_*1*_ (Table 2), its *α’* value may be closer to 1 than estimated. Bonobos are not expected to have ERO males given their strongly matriarchal society [8, 10, 49]. It is possible that their *V*_*M*_*(K)* is unusually small whereas *V*_*F*_*(K)* is unusually large, thus giving rise to *α’* ≤ 1.

Gorillas have the second smallest *α’* that falls between 1 and 10. They usually have a harem structure [9, 50-52] whereby *α’* is expected to be > 1. However, *α’* may not be very high since males do not have access to a large number of females to become ERO males. For chimpanzees, humans, and Sumatran orangutans, *α’* values are likely ≥ 20, reflecting a very large *V*_*M*_*(K)*. While both orangutan species share a semi-solitary lifestyle, Sumatran orangutans live in higher-density populations with more frequent social interactions, increased male competition, and greater access to females than Bornean orangutan [61-63]. This heightened competition may lead to greater reproductive skew, which explains the higher *α’* values observed in Sumatran orangutans.

While this study focuses on humans and great apes, the reported patterns may be general. Table 3 compiles field observations of 3 species of old-world monkeys and one rodent species. Although the reported K*’s are observed progeny numbers rather than the formulated K (reproductive success), the trend is as expected. The ratio of progeny variance between males and females is indeed much larger in species of polygeny than in species of polyandry or monogamy. However, field observations do not yield *α’* estimates, which are obtainable only from the genomic data as explained below.

**Table 3.**
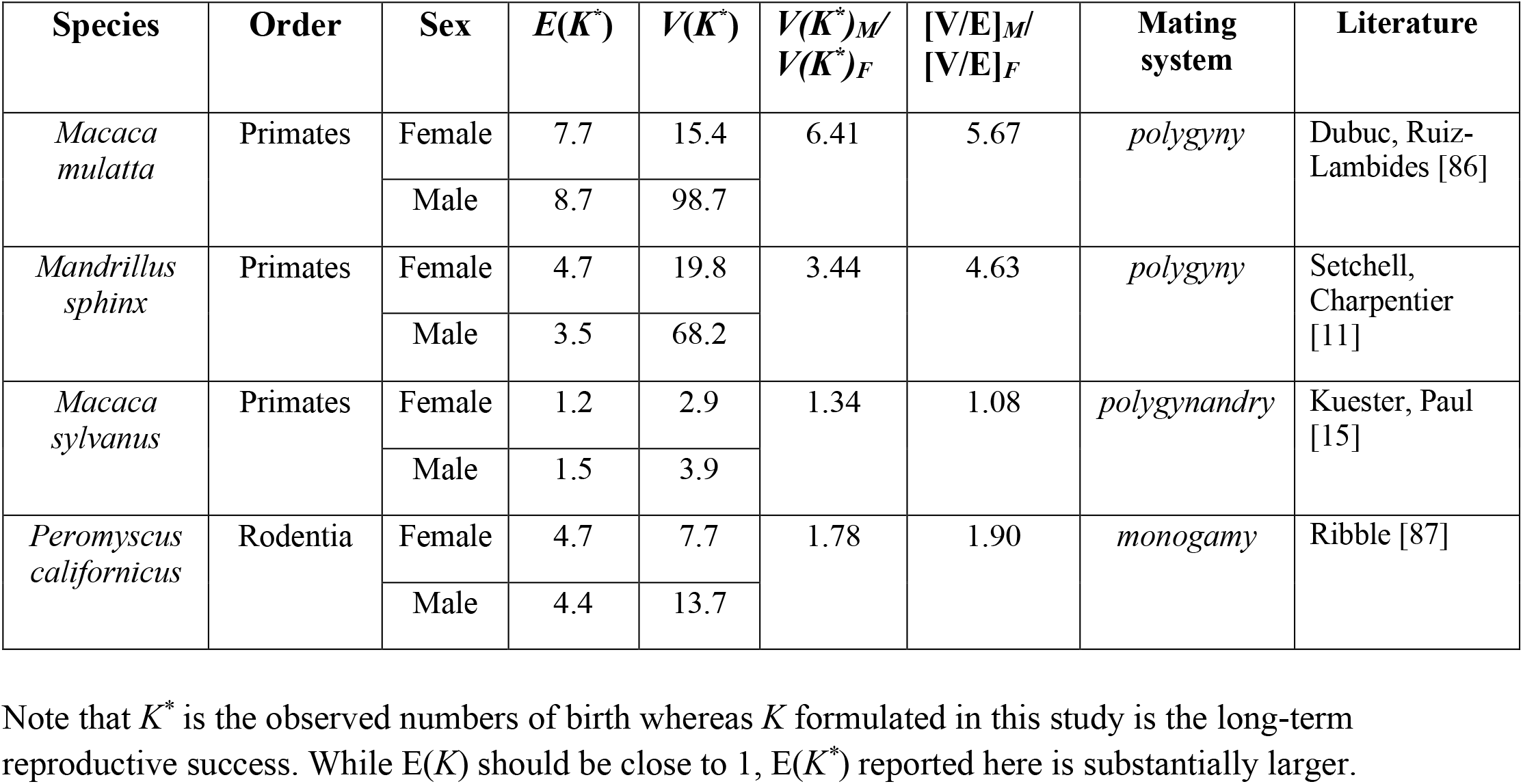
Field observations of progeny numbers (*K*^*\**^) in non-ape mammals under three different socio-sexual structures.

The high *α’* values so obtained for humans and great apes are interpreted to mean a small percentage of ERO males. Note that a high *α’* is equivalent to a high *V*_*M*_*(K)* which can be due to either a few outlier males with very large breeding successes or many males with moderate successes. The former is favored because of the constraint of *E*_*M*_*(K)* = *E*_*F*_*(K)* ∼ 1 [= *E(K)*] as *E(K)* should be close to 1 over the evolutionary time scale. Thus, if many males have high reproductive successes, females should be equally productive, thus raising *E(K)* to an unsustainable level. For example, if 10% of males produce *K* = 5 progeny (while other males and all females having *E(K)* = *V(K)* = 1), *α’* would be < 2, but *E(K)* would be nearly 1.5. On the other hand, if there are only 1% outlier males with *K* = 40, then *α’* would be > 15 with a similar *E(K)*, while allowing some males to have no offspring can maintain *E(K)* close to 1. In other words, only large *K*’s can lead to large *α’* without substantially increasing *E(K)*. In Supplementary Note 4 and Fig. S2, we provide a more formal analysis using the Gamma distribution.

It is noteworthy that the long-term stochastic effect is determined by the generation with the largest variation. (Thus, the long-term *N*_*e*_ is the harmonic mean of *N*’s across generations.) Therefore, ERO males only need to emerge once in a while to have the large impact on evolution.

1. Social-sexual conditions - Many factors of the social-sexual structure may contribute to the emergence of ERO males and large *α’* values. We shall discuss only two such factors. First, the absence of paternal care may permit males to pursue ever higher *K*’s. That may be part of the differences between chimpanzees and gorillas, as gorilla fathers are known to provide protection [12], while paternal care in chimpanzees is rare or negligible [64]. Neither do orangutans show paternal care [12, 65]. For example, ERO males have been observed in *Macaca mulatta* and *Mandrillus sphinx*. Both males provide little paternal care and can sire a maximum of 47 and 41 offspring [11, 14], respectively, much higher than the reproductive output of females.

Second, and most interesting, the reproductive advantage of males can be amplified *non-genetically*. Powerful parents, either maternal or paternal, can help their sons gain social-sexual advantages [66, 67]. So do successful brothers who form coalitions [8]. Consequently, the advantages in one generation may be amplified in several subsequent generations, as speculated [68]. Humans and chimpanzees share some of these behavioral traits. They include male philopatry with female-biased dispersal [69, 70], allowing males to remain in their natal groups and form male–male cooperation within strong alliances [8, 71, 72]. Furthermore, maternal support for sons in their pursuit of social dominance and mating opportunities are also well known [8, 71, 72]. Genomic studies have indeed suggested that humans are closer to chimpanzees than to gorillas in their male reproductive strategies [2]. A new direction taken by neuro-scientists who use non-human primates as a research model may be promising in resolving some of these issues[73-76]. Field studies such as those on colobine monkeys [77] should be able to discover additional social conditions for the emergence of ERO males.

iii)Other tests – It should be noted that the ERO phenomenon reflects the average over the evolutionary time span, close to the coalescence time of Y-linked genes, roughly 50-250 thousand years in humans[4]. In contrast, observations in field studies reflect social-sexual patterns in specific ethnic group during a specific period. The two lines of evidence are not always compatible. For example, in Brown et al.’s survey[31] of modern human society, *α’* is between 1 and 5. A possible explanation can be found in the studies of Yanomama and other South American ethnic group. J. V. Neel [5] observed that the unacculturated ethnic group tend to have a few ERO males (or headmen) that sire large number of children and grandchildren. Such strong reproductive biases may also account for the unusually large genetic differentiations among local tribes. In this context, there would be particular interest in humans’ recent past[13, 78, 79], whereby “field observations” in the form of historical documents and sociological records are extensive. Hence, if we can infer the recently accumulated variations on the A, X and Y chromosomes [80, 81], field observations and genomic data would be directly comparable.

In conclusion, while previous studies have invoked either selection or sex differences in breeding success (but not both [4, 21, 22, 38]), this study takes into account both forces (see Supplementary Note 5 for discussions on positive vs. negative selection). Consequently, the estimation of *α’* reveals an extreme form of sexual selection that could only be speculated before.

## Materials and Methods

### Non-human Data collection, *θ*_*w*_ estimation and human data processing

Details are presented in Supplement

### The *α’* estimate with the minimum MSE (Mean Squared Error)

To mitigate the issue of the *α’* estimation’s extreme sensitivity to the measurement errors in chromosome diversity and mutation rate, we employed a combination of Eq. (1-3) and minimum MSE to determine the optimal *α’*. We generated a range of potential (α^*^) values ranging from 0 to 200, with increments of 0.001. For each *α’*, we computed the expected three diversity ratios Exp[R_YA_], Exp[R_XA_], and Exp[R_YX_], using Equations in Supplementary Note 1. Subsequently, based on the observed diversity ratios, Obs[R_YA_], Obs[R_XA_], and Obs[R_YX_],and the calculated Exp[R], we estimated the MSE for each *α’* as follows:

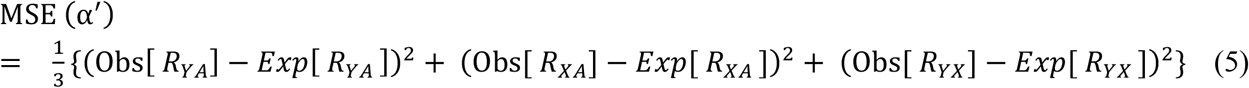

The *α’* value corresponding to the minimum MSE was used in Tables 1 and 2. Values exceeding 200 are designated as “Inf”. We applied this method to investigate the *α’* estimate with the minimum MSE, incorporating three observed diversity ratios for each pair of chromosomes.

### Bootstrapping procedure

Resampling DNA segments – The noncoding portion of chromosome sequences from 84 individuals in the HGDP dataset were grouped by chromosome type (n= 84 for X, Y, and autosomes) and aligned within each group. The pooled alignments were then divided into non-overlapping 1-kb segments, yielding 2,875,310 segments for the haploid autosomes, 156,060 for the X and 57,240 for the Y. Bootstrapping was re-iterated 10,000 times. Each time, 120,000 segments were sampled with replacement from autosomes, X and Y for the calculation of *α’*, based on the resampled *θ*_*w*_ and *θ*_*1*_, respectively (see Supporting Materials for details). Note that *α’* ± Dev1 in Table 2 covers ≥ 95% of the bootstrapping values.

Resampling individuals – In each of the 100 iterations, 84 individuals were sampled with replacement from the HGDP dataset (n = 84). For each bootstrap iteration, *α’* was estimated based on *θ*_*w*_ and *θ*_*1*_, following the same procedure as applied to the original dataset. The reported *α’* ± Dev2 in Table 2 represents the interval covering ≥ 95% of the bootstrapping values.

### Linear extrapolation for F(*x* ∼ 0)

Here, Eq. (6) is an extension of [82], and in a somewhat different form from [83], for the effect of selection on the site frequency spectrum (SFS) (detailed derivations given in Supplementary Note 2),

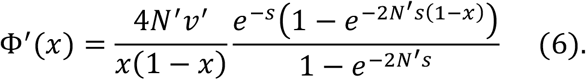

When *s* = 0, Eq. (6) is reduced to

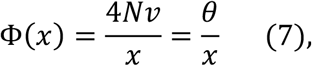

as is generally known. Φ(*x*) is the probability density of mutations of frequency *x* in the population. Here, we use Eq. (6) to portray the mutation spectrum of Y-linked mutations under selection, denoting the variant frequency, population size, and mutation rate as *x’, N’* and *v’*, respectively. The simpler Eq. (7) is for autosomal mutations. What we need is the SFS ratio of Y to Autosome, hence,

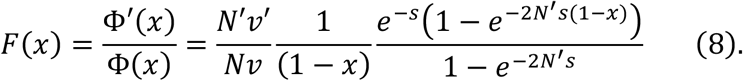

While *s* can be positive or negative selection, the numerical results presented will only be about cases of *s* < 0; i.e., the so-called background selection on Y [22, 38, 84]. *N* is for the autosomes. For the Y chromosome, *N’* = *ηN* where *η* = 0.5 in the standard WF population as only half of the population has Y. Importantly, *η* < 0.5 is a measure of *V*_*M*_*(K)*. Furthermore, because *N* is at least 1000 and *e* ^−,*N ηs*^ ≫ 1. F(*x*) in the logarithmic form when *x* << 1 would be Eq. (4), with further details provided in Supplementary Note 3.

To estimate the F(*x* ∼ 0) through linear extrapolation, we utilized *ξ*_*i*_ in human population (Table S3), where *i* = 1, 2, 3, 4. Based on Eq. (4), we defined the ratio of F(0) to F(1/*n*) as *Ω*,

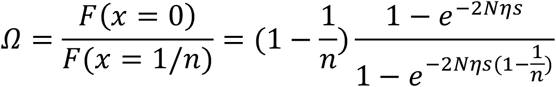

Considering the *N* is large and *s* is a negative value of deleterious mutations, both 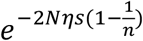 and *e*^−,*N ηs*^ ≫ 1. Therefore, with transformed,

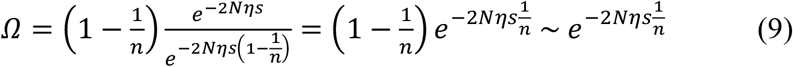

We set *N* = 7500, with sample sizes of *n* = 84 for HGDP and *n* = 46 for KGP. *Ω* was simplified as *e*^−,179*ηs*^ and *e*^−,326 *ηs*^ accordingly.

For each dataset, we generated pairs of ln[F(*i*/*n*)] and ln[F(*j*/*n*)], where 0 ≤ *i* < *j* and *j* ≤ 4, resulting in 10 slopes for five data points. Notably, the slope between F*(*x* ∼ 0) and F(1/*n*) was treated as a dummy number and was not used in the analysis. Next, we employed the 9 slopes generated from Eq. (4) along with Eq. (9) to produce a set of *Ω*. Each slope was used to calculate a distinct F(*x* ∼ 0) estimate, thus yielding 9 different *α’* estimates from one dataset.

Finally, we extrapolated F(*x* ∼ 0) from F(1/*n*)

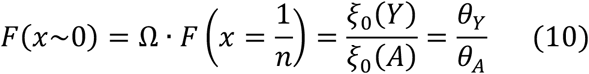

Hence, we obtained *R*_*YA*_ = F(*x* ∼ 0). By applying *R*_*YA*_ to Eq. (1), we derived 9 *α’*(*YA*) values for each population from one dataset (Table S4).

## References

1. Short RV. Sexual Selection and Its Component Parts, Somatic and Genital Selection, as Illustrated by Man and the Great Apes. In: Rosenblatt JS, Hinde RA, Beer C, et al. (eds.). Advances in the Study of Behavior: Academic Press; 1979. 131–158.

2. Wyckoff GJ, Wang W, Wu C-I. Rapid evolution of male reproductive genes in the descent of man. Nature. 2000; 403(6767): 304–309. doi: 10.1038/35002070

3. Dixson AF. Sexual selection and the origins of human mating systems: Oxford University Press, USA, 2009.

4. Karmin M, Saag L, Vicente M et al. A recent bottleneck of Y chromosome diversity coincides with a global change in culture. Genome Res. 2015; 25(4): 459–466. doi: 10.1101/gr.186684.114

5. Neel JV. Lessons from a “Primitive” People. Science (New York, NY). 1970; 170(3960): 815–822. doi: 10.1126/science.170.3960.815

6. Wade Michael J, Shuster Stephen M. Sexual Selection: Harem Size and the Variance in Male Reproductive Success. The American Naturalist. 2004; 164(4): E83–E89. doi: 10.1086/424531

7. Wroblewski EE, Murray CM, Keele BF et al. Male dominance rank and reproductive success in chimpanzees, Pan troglodytes schweinfurthii. Animal Behaviour. 2009; 77(4): 873–885. doi: 10.1016/j.anbehav.2008.12.014

8. Gruber T, Clay Z. A Comparison Between Bonobos and Chimpanzees: A Review and Update. Evolutionary Anthropology: Issues, News, and Reviews. 2016; 25(5): 239–252. doi: 10.1002/evan.21501

9. McGrew WC, Marchant LF, Nishida T. Great ape societies: Cambridge University Press, 1996.

10. De Waal FB, Lanting F. Bonobo: The forgotten ape: Univ of California Press, 2023.

11. Setchell JM, Charpentier M, Wickings EJ. Sexual selection and reproductive careers in mandrills (Mandrillus sphinx). Behavioral Ecology and Sociobiology. 2005; 58(5): 474–485. doi: 10.1007/s00265-005-0946-2

12. Smuts BB, Cheney DL, Seyfarth RM et al. Primate societies: University of Chicago Press, 2008.

13. Ning Z, Tan X, Yuan Y et al. Expression profiles of east–west highly differentiated genes in Uyghur genomes. National Science Review. 2023; 10(4): nwad077. doi: 10.1093/nsr/nwad077

14. Dubuc C, Ruiz-Lambides A, Widdig A. Variance in male lifetime reproductive success and estimation of the degree of polygyny in a primate. Behavioral Ecology. 2014; 25(4): 878–889. doi: 10.1093/beheco/aru052

15. Kuester J, Paul A, Arnemann J. Age-related and individual differences of reproductive success in male and female barbary macaques (Macaca sylvanus). Primates. 1995; 36(4): 461–476. doi: 10.1007/BF02382869

16. Balaresque P, Poulet N, Cussat-Blanc S et al. Y-chromosome descent clusters and male differential reproductive success: young lineage expansions dominate Asian pastoral nomadic populations. European Journal of Human Genetics. 2015; 23(10): 1413–1422. doi: 10.1038/ejhg.2014.285

17. Jobling MA, Tyler-Smith C. Human Y-chromosome variation in the genome-sequencing era. Nature Reviews Genetics. 2017; 18(8): 485–497. doi: 10.1038/nrg.2017.36

18. Xue Y, Zerjal T, Bao W et al. Recent spread of a Y-chromosomal lineage in northern China and Mongolia. American journal of human genetics. 2005; 77(6): 1112–1116. doi: 10.1086/498583

19. Miyata T, Hayashida H, Kuma K et al. Male-driven molecular evolution: a model and nucleotide sequence analysis. Cold Spring Harbor symposia on quantitative biology. 1987; 52: 863–867. doi: 10.1101/sqb.1987.052.01.094

20. Rozen S, Marszalek JD, Alagappan RK et al. Remarkably little variation in proteins encoded by the Y chromosome’s single-copy genes, implying effective purifying selection. American journal of human genetics. 2009; 85(6): 923–928. doi: 10.1016/j.ajhg.2009.11.011

21. Gerrard DT, Filatov DA. Positive and negative selection on mammalian Y chromosomes. Molecular Biology and Evolution. 2005; 22(6): 1423–1432. doi: DOI 10.1093/molbev/msi128

22. Wilson Sayres MA, Lohmueller KE, Nielsen R. Natural selection reduced diversity on human y chromosomes. Plos Genet. 2014; 10(1): e1004064. doi: 10.1371/journal.pgen.1004064

23. Charlesworth B. The effect of life-history and mode of inheritance on neutral genetic variability. Genet Res. 2001; 77(2): 153–166. doi: 10.1017/s0016672301004979

24. Wilder JA, Mobasher Z, Hammer MF. Genetic evidence for unequal effective population sizes of human females and males. Mol Biol Evol. 2004; 21(11): 2047–2057. doi: 10.1093/molbev/msh214

25. Nam K, Munch K, Hobolth A et al. Extreme selective sweeps independently targeted the X chromosomes of the great apes. Proceedings of the National Academy of Sciences. 2015; 112(20): 6413–6418. doi: 10.1073/pnas.1419306112

26. Charlesworth B, Charlesworth D. The degeneration of Y chromosomes. Philosophical transactions of the Royal Society of London Series B, Biological sciences. 2000; 355(1403): 1563–1572. doi: 10.1098/rstb.2000.0717

27. Xu J, Peng X, Chen Y et al. Free-living human cells reconfigure their chromosomes in the evolution back to uni-cellularity. eLife. 2017; 6. doi: 10.7554/eLife.28070

28. Gong G, Xiong Y, Xiao S et al. Origin and chromatin remodeling of young X/Y sex chromosomes in catfish with sexual plasticity. National Science Review. 2023; 10(2): nwac239. doi: 10.1093/nsr/nwac239

29. Watterson GA. On the number of segregating sites in genetical models without recombination. Theoretical Population Biology. 1975; 7(2): 256–276. doi: 10.1016/0040-5809(75)90020-9

30. Hallast P, Maisano Delser P, Batini C et al. Great ape Y Chromosome and mitochondrial DNA phylogenies reflect subspecies structure and patterns of mating and dispersal. Genome Res. 2016; 26(4): 427–439. doi: 10.1101/gr.198754.115

31. Brown GR, Laland KN, Mulder MB. Bateman’s principles and human sex roles. Trends in Ecology & Evolution. 2009; 24(6): 297–304. doi: 10.1016/j.tree.2009.02.005

32. Kimura M, Crow JF. The Measurement of Effective Population Number. Evolution. 1963; 17(3): 279–288. doi: 10.1111/j.1558-5646.1963.tb03281.x

33. Crow JF, Kimura M. An Introduction to Population Genetics Theory: Blackburn Press, 1970.

34. Chen Y, Tong D, Wu CI. A New Formulation of Random Genetic Drift and Its Application to the Evolution of Cell Populations. Mol Biol Evol. 2017; 34(8): 2057–2064. doi: 10.1093/molbev/msx161

35. Hou M, Shi JR, Gong ZK et al. Intra-vs. Interhost Evolution of SARS-CoV-2 Driven by Uncorrelated Selection-The Evolution Thwarted. Molecular Biology and Evolution. 2023; 40(9): msad204. doi: ARTN msad204 10.1093/molbev/msad204

36. Fay JC, Wu CI. Hitchhiking under positive Darwinian selection. Genetics. 2000; 155(3): 1405–1413. doi: 10.1093/genetics/155.3.1405

37. Fu Y-X. Statistical Tests of Neutrality of Mutations Against Population Growth Hitchhiking and Background Selection. Genetics. 1997; 147(2): 915–925. doi: 10.1093/genetics/147.2.915

38. Charlesworth B, Morgan MT, Charlesworth D. The effect of deleterious mutations on neutral molecular variation. Genetics. 1993; 134(4): 1289–1303. doi: 10.1093/genetics/134.4.1289

39. Makova KD, Li WH. Strong male-driven evolution of DNA sequences in humans and apes. Nature. 2002; 416(6881): 624–626. doi: 10.1038/416624a

40. Wu FL, Strand AI, Cox LA et al. A comparison of humans and baboons suggests germline mutation rates do not track cell divisions. PLoS Biol. 2020; 18(8): e3000838. doi: 10.1371/journal.pbio.3000838

41. Li W-H. Molecular evolution. Sunderland, Mass.: Sinauer Associates, 1997.

42. Huang W, Chang BH, Gu X et al. Sex differences in mutation rate in higher primates estimated from AMG intron sequences. J Mol Evol. 1997; 44(4): 463–465. doi: 10.1007/pl00006166

43. Bachtrog D. Y-chromosome evolution: emerging insights into processes of Y-chromosome degeneration. Nature Reviews Genetics. 2013; 14(2): 113–124. doi: 10.1038/nrg3366

44. Betancourt AJ, Presgraves DC. Linkage limits the power of natural selection in Drosophila. Proceedings of the National Academy of Sciences. 2002; 99(21): 13616–13620. doi: 10.1073/pnas.212277199

45. Tajima F. Statistical method for testing the neutral mutation hypothesis by DNA polymorphism. Genetics. 1989; 123(3): 585–595. doi: 10.1093/genetics/123.3.585

46. Fu YX, Li WH. Statistical tests of neutrality of mutations. Genetics. 1993; 133(3): 693–709. doi: 10.1093/genetics/133.3.693

47. Arbiza L, Gottipati S, Siepel A et al. Contrasting X-linked and autosomal diversity across 14 human populations. American journal of human genetics. 2014; 94(6): 827–844. doi: 10.1016/j.ajhg.2014.04.011

48. Xue Y, Prado-Martinez J, Sudmant PH et al. Mountain gorilla genomes reveal the impact of long-term population decline and inbreeding. Science (New York, NY). 2015; 348(6231): 242–245. doi: 10.1126/science.aaa3952

49. Furuichi T. Female Contributions to the Peaceful Nature of Bonobo Society. Evol Anthropol. 2011; 20(4): 131–142. doi: 10.1002/evan.20308

50. Harcourt AH, Stewart KJ. Gorilla society: What we know and don’t know. Evol Anthropol. 2007; 16(4): 147–158. doi: 10.1002/evan.20142

51. Doran DM, McNeilage A. Gorilla ecology and behavior. Evolutionary Anthropology: Issues, News, and Reviews. 1998; 6(4): 120–131. doi: 10.1002/(SICI)1520-6505(1998)6:4<120::AID-EVAN2>3.0.CO;2-H

52. Watts DP. Mountain gorilla reproduction and sexual behavior. American Journal of Primatology. 1991; 24(3-4): 211–225. doi: 10.1002/ajp.1350240307

53. Bergström A, McCarthy SA, Hui R et al. Insights into human genetic variation and population history from 929 diverse genomes. Science (New York, NY). 2020; 367(6484). doi: 10.1126/science.aay5012

54. Abecasis GR, Altshuler D, Auton A et al. A map of human genome variation from population-scale sequencing. Nature. 2010; 467(7319): 1061–1073. doi: 10.1038/nature09534

55. Bradley E. Second Thoughts on the Bootstrap. Statistical Science. 2003; 18(2): 135–140. doi: 10.1214/ss/1063994968

56. Efron B. Bootstrap Methods: Another Look at the Jackknife. The Annals of Statistics. 1979; 7(1): 1–26. doi: 10.1214/aos/1176344552

57. Takahata N. Allelic genealogy and human evolution. Mol Biol Evol. 1993; 10(1): 2–22. doi: 10.1093/oxfordjournals.molbev.a039995

58. Tenesa A, Navarro P, Hayes BJ et al. Recent human effective population size estimated from linkage disequilibrium. Genome Res. 2007; 17(4): 520–526. doi: 10.1101/gr.6023607

59. Irwin DE. Sex chromosomes and speciation in birds and other ZW systems. Molecular ecology. 2018; 27(19): 3831–3851. doi: 10.1111/mec.14537

60. Smeds L, Warmuth V, Bolivar P et al. Evolutionary analysis of the female-specific avian W chromosome. Nature Communications. 2015; 6(1): 7330. doi: 10.1038/ncomms8330

61. Kunz JA, Duvot GJ, van Noordwijk MA et al. The cost of associating with males for Bornean and Sumatran female orangutans: a hidden form of sexual conflict? Behavioral Ecology and Sociobiology. 2020; 75(1): 6. doi: 10.1007/s00265-020-02948-4

62. Mitra Setia T, Delgado RA, Utami Atmoko SS et al. 245Social organization and male–female relationships. Orangutans: Geographic Variation in Behavioral Ecology and Conservation: Oxford University Press; 2008. 0.

63. Utami Atmoko SS, Mitra Setia T, Goossens B et al. 235Orangutan mating behavior and strategies. Orangutans: Geographic Variation in Behavioral Ecology and Conservation: Oxford University Press; 2008. 0.

64. Nishida T. Chimpanzees of the Mahale Mountains 1990.

65. Allman J, Rosin A, Kumar R et al. Parenting and survival in anthropoid primates: Caretakers live longer. Proceedings of the National Academy of Sciences. 1998; 95(12): 6866–6869. doi: 10.1073/pnas.95.12.6866

66. Surbeck M, Boesch C, Crockford C et al. Males with a mother living in their group have higher paternity success in bonobos but not chimpanzees. Curr Biol. 2019; 29(10): R354–R355. doi: 10.1016/j.cub.2019.03.040

67. Alberts SC, Altmann J. Preparation and activation: determinants of age at reproductive maturity in male baboons. Behavioral Ecology and Sociobiology. 1995; 36(6): 397–406. doi: 10.1007/BF00177335

68. Zerjal T, Xue Y, Bertorelle G et al. The genetic legacy of the Mongols. The American Journal of Human Genetics. 2003; 72(3): 717–721.

69. Goodall J. The chimpanzees of Gombe: patterns of behavior. : Cambridge: Harvard University Press, 1986.

70. Kano T. The last ape: pygmy chimpanzee behavior and ecology: Stanford University Press Stanford, 1992.

71. Nishida T. Chimpanzees of the lakeshore: natural history and culture at Mahale: Cambridge University Press, 2011.

72. Boesch C. The real chimpanzee: sex strategies in the forest: Cambridge University Press, 2009.

73. Poo M-m. Editorial of non-human primate research. National Science Review. 2023; 10(11): nwad326. doi: 10.1093/nsr/nwad326

74. Liu X, Cheng Z, Lin H et al. Decoding effects of psychoactive drugs in a high-dimensional space of eye movements in monkeys. National Science Review. 2023; 10(11): nwad255. doi: 10.1093/nsr/nwad255

75. Meng X, Lin Q, Zeng X et al. Brain developmental and cortical connectivity changes in transgenic monkeys carrying the human-specific duplicated gene SRGAP2C. National Science Review. 2023; 10(11): nwad281. doi: 10.1093/nsr/nwad281

76. Zhang R, Quan H, Wang Y et al. Neurogenesis in primates versus rodents and the value of non-human primate models. National Science Review. 2023; 10(11): nwad248. doi: 10.1093/nsr/nwad248

77. Qi X-G, Wu J, Zhao L et al. Adaptations to a cold climate promoted social evolution in Asian colobine primates. Science (New York, NY). 380(6648): eabl8621. doi: 10.1126/science.abl8621

78. Wen J, Liu J, Feng Q et al. Ancestral origins and post-admixture adaptive evolution of highland Tajiks. National Science Review. 2024; 11(9): nwae284. doi: 10.1093/nsr/nwae284

79. Wu C-I. The genetics of race differentiation—should it be studied? National Science Review. 2023; 10(4): nwad068. doi: 10.1093/nsr/nwad068

80. Li H, Durbin R. Inference of human population history from individual whole-genome sequences. Nature. 2011; 475(7357): 493–496. doi: 10.1038/nature10231

81. Liu X, Fu Y-X. Exploring population size changes using SNP frequency spectra. Nature genetics. 2015; 47(5): 555–559. doi: 10.1038/ng.3254

82. Kimura M. The Number of Heterozygous Nucleotide Sites Maintained in a Finite Population Due to Steady Flux of Mutations. Genetics. 1969; 61(4): 893–903. doi: 10.1093/genetics/61.4.893

83. Ewens WJ. The sampling theory of selectively neutral alleles. Theoretical Population Biology. 1972; 3(1): 87–112. doi: 10.1016/0040-5809(72)90035-4

84. Charlesworth B. The effects of deleterious mutations on evolution at linked sites. Genetics. 2012; 190(1): 5–22. doi: 10.1534/genetics.111.134288

85. Stone AC, Griffiths RC, Zegura SL et al. High levels of Y-chromosome nucleotide diversity in the genus Pan. Proceedings of the National Academy of Sciences. 2002; 99(1): 43–48. doi: 10.1073/pnas.012364999

86. Dubuc C, Ruiz-Lambides A, Widdig A. Variance in male lifetime reproductive success and estimation of the degree of polygyny in a primate. Behav Ecol. 2014; 25(4): 878–889. doi: 10.1093/beheco/aru052

87. Ribble DO. Lifetime Reproductive Success and its Correlates in the Monogamous Rodent, Peromyscus californicus. The Journal of Animal Ecology. 1992; 61(2): 457–468. doi: 10.2307/5336

