## supplementary information for "Genomic Evidence for Males of Exceptional Reproductive Output (ERO) in Apes and Humans"

**This PDF file includes:**

Supporting Materials
Supporting text
Figures S1 to S2
Tables S1, S2, S4
Legends for Table S3, S5
SI References

**Other supporting materials for this manuscript include the following:**

Table S3
Table S5

### Supporting Materials

#### Non-human Data collection and $\theta_w$ estimation

Data on each chromosome's diversity in non-human great apes was gathered from published studies. Y and X chromosomes data were collected by [1] with a total sample of 4 bonobos, 19 chimpanzees (subspecies: 9 Western, 3 Eastern, 4 Nigeria-Cameroon, and 3 Central chimpanzees), 14 gorillas (subspecies: 7 Western lowland, 3 Eastern lowland, and 3 Mountain gorillas) and 6 Orangutans (subspecies: 4 Sumatran and 2 Bornean). The male-specific region of the Y chromosome (MSY) was based on a human-reference-sequence, excluding the ampliconic and X-transposed regions, resulting in 2.61 and 3.80 Mb (depending on species) for analysis. In this data set, the 3 central chimpanzees (*Pan troglodytes troglodytes*), exhibited > 10-fold higher diversity compared to other chimpanzee populations and were identified as belonging to divergent populations, thus excluded from the analysis. The autosomal diversity of non-human great apes (utilizing only chromosome 1) was sourced from [2], with the exception of the Gorilla genus obtained from [3].

The genetic variation of A (autosome), Y, and X chromosomes in non-human great apes is summarized in Table S1. Genetic diversity $\theta_w$ estimated by counting the number of segregating sites (S) using the Watterson estimator [4].

#### Human data processing

$\theta_w$ and $\theta_\pi$ values for each chromosome of human, covering both coding and non-coding regions, were computed using data from Human Genome Diversity Project (HGDP) [5] and 1000 Genomes Project (KGP) [6] across three major populations: African, European, and Asian (Table S2). The sample sizes were 84 and 46 for each population in the two data sets, respectively (sample list shown in Table S5). The coding sequences (CDS) of genes was retained as the coding region, while the complementary region was considered non-coding from https://ftp.ensembl.org/pub/release-111/gff3/homo_sapiens.

To reduce the effects of miscalling, we only included variants that passed the 1000 Genomes Project's strict mask (20160622 version). Furthermore, to ensure consistency with released Phase 3 of the 1000 Genomes Project, we performed statistical phasing to the HGDP dataset. We used SHAPEIT4 [7] and Eagle2 [8] with default parameter settings for the phasing of high-quality SNPs on autosomes and X chromosome. For the analysis of chromosome Y, loci with >10% genotype missingness were excluded, and only variants within the male-specific region of the Y chromosome (MSY) were retained, resulting in 13.9 Mb for downstream analysis. This yielded the final set of 77,046 variants for HGDP, and 59,386 variants for KGP in MSY.

After haplotype phasing per chromosome, we presented the number of sites, $\xi_i$ (= $\theta/i$), with the variant occurring $i$ times in each population, with $i$ = 1, 2, 3, 4, 5 (Table S3). Since $\xi_i$ may be frequently inflated by sequencing errors, we used this statistic only on the high-quality data that have been carefully checked for this purpose, ensuring, for example, each singleton was supported by at least 5 mapped reads. To reduce estimation bias, we used in-house scripts to perform 10,000 bootstrap iterations. In each iteration, 120,000 1-kb windows were randomly sampled from the genome (including the X, Y, and autosomes), and the mean values and 95% confidence intervals (CIs) were calculated.

The Y chromosome, excluding the pseudoautosomal regions, is fully linked. The accuracy of the $\theta$ estimation with or without recombination is addressed by diffusion theory.

We will use only $\theta_W$ estimation for illustration but the logic is the same for other theta estimates such as $\theta_\pi$. The observed quantity for $\theta_W$ estimation is $W$ [9].

In the absence of recombination, the mean and variance of $W$ are [4, 10]

$E(W) = \theta$

$V(W) = \theta/a_n + b_n \theta^2/ a_n^2$

where $n$ is the number of sequences, $a_n = 1+ \frac{1}{2} + \ldots + 1/(n-1)$ and $b_n = 1+ \frac{1}{4} + \ldots + 1/(n-1)^2$.

With free recombination ($r = 0.5$ among all variants, the extreme opposite):

$E(W) = \theta$

$V(W) = \theta/a_n$

It can be seen that recombination does not affect E($W$), but it reduces V($W$). It is true that the variance of the estimate of $\theta$ without recombination would be larger. However, even if we assume that X and autosomal variants are freely recombining (which is an exaggeration), the difference in V($W$) would not be large mainly because $\theta$ is rather small on the Y chromosome. By the coalescence interpretation, small $\theta$ means low level of standing variation and the influence of recombination would be diminished. We can also use the short sojourn time in the population to bolster the argument.

We are primarily interested in very low frequency variants because this part of SFS is relatively unaffected by selection. In addition, low-frequency SFSs are also less affected by recombination, as explained above.

#### Supporting Information Text

**Note 1. Estimation of the ratio of male-to-female reproductive output, *α'***

Let $K$ be the progeny number of each individual and the mean value of $K$ for males and females is $E_M(K)$ and $E_F(K)$, respectively. The variance of $K$ is hence $V_M(K)$ and $V_F(K)$. Clearly, $E_M(K) = E_F(K)$ if the sex ratio is 1:1.

The male-to-female ratio of $K$ variance, $V_M(K)/V_F(K)$, referred to as $\alpha'$, is of interest. ($\alpha'$ is used because $\alpha$ (without the prime) has been commonly used for the male-to-female ratio in mutation rate.)) Thus,

$$\alpha' = \frac{V_M(K)}{V_F(K)}$$

During evolution, since Y, X, and A (autosome) spend 100%, 1/3, and 1/2 of the time in males and the rest in females,

$$V_A(K) = \frac{1}{2}V_F(K) + \frac{1}{2}V_M(K)$$

$$V_X(K) = \frac{2}{3}V_F(K) + \frac{1}{3}V_M(K)$$

$$V_Y(K) = V_M(K)$$

We then obtain the relationship between the $K$ variance and $\alpha'$ as

$$\frac{V_Y(K)}{V_A(K)} = \frac{2\alpha'}{1+\alpha'}$$

$$\frac{V_X(K)}{V_A(K)} = \frac{2}{3} \cdot \frac{2+\alpha'}{1+\alpha'}$$

$$\frac{V_Y(K)}{V_X(K)} = \frac{3\alpha'}{2+\alpha'}$$

The mutation rates for X and Y chromosomes are denoted as $x\mu$ and $y\mu$, where $\mu$ represents the mutation rate for autosomes. Combining with the $\theta_A$, $\theta_X$, or $\theta_Y$ ($\theta$ can be estimated from polymorphic DNA sequences), we have these three ratios:

$$R_{YA} = \frac{\theta_Y}{\theta_A}$$

$$R_{XA} = \frac{\theta_X}{\theta_A}$$

$$R_{YX} = \frac{\theta_Y}{\theta_X}$$

where $\theta$ is equal to $N_e\mu$.

We note that $N_e = N/V(K)$ and the relative values of $N$'s for Y, X, and A are 1:3:4. The relative values of $\mu$'s for Y, X, and A are $y$: $x$: 1. In equilibrium populations, we obtain

$$R_{YA} = \frac{\frac{N(Y)}{V_Y(K)} \cdot yu}{\frac{N(A)}{V_A(K)} \cdot u}; \; R_{XA} = \frac{\frac{N(X)}{V_X(K)} \cdot xu}{\frac{N(A)}{V_A(K)} \cdot u}; \; R_{YX} = \frac{\frac{N(Y)}{V_Y(K)} \cdot yu}{\frac{N(X)}{V_X(K)} \cdot xu}.$$

which can be simplified to

$$R_{YA} = \frac{y}{4} \cdot \frac{1+\alpha'}{2\alpha'}$$

$$R_{XA} = \frac{9x}{8} \cdot \frac{1+\alpha'}{2+\alpha'}$$

$$R_{YX} = \frac{y}{3x} \cdot \frac{2+\alpha'}{3\alpha'}$$

The *α'* is then:

$$\alpha'(YA) = \frac{y}{8R_{YA} - y}$$

$$\alpha'(XA) = \frac{16R_{XA} - 9x}{9x - 8R_{XA}}$$

$$\alpha'(YX) = \frac{2y}{9xR_{YX} - y}$$

**Note 2. Site frequency spectrum (SFS) under selection**

To obtain the SFS under selection, we have to use the diffusion equations, in which the mean and variance of the rate of change in frequency *x* is the key function.

$$E(\delta x) = M_\delta(x)\delta t + o(\delta t) \ (1)$$

$$V(\delta x) = V_\delta(x)\delta t + o(\delta t) \ (2)$$

We will consider a more general case (diploid population here) in which there is arbitrary degree of dominance. i.e., the selective advantages (for deleterious mutation, $s < 0$) of $A_1A_1$, and $A_1A_2$ over $A_2A_2$ are *s* and *sh* respectively. Let us assume a pair of alleles $A_1$ and $A_2$ with respective frequencies *x* and 1-*x*. And let $u_{12}$ be the mutation rate from $A_1$ to $A_2$ and let $u_{21}$ be the mutation rate in the reverse. According to Eq. (9.3.1) in [11],

$$M_\delta(x) = -u_{12}x + u_{21}(1-x) + sx(1-x)[h + (1-2h)x] \ (3)$$

$$V_\delta(x) = \frac{x(1-x)}{2N_e} \quad (4)$$

According to Eq. (37) in [12], the site frequency spectrum at steady-state is

$$\Phi(x) = \frac{2v}{V_\delta(x)G(x)} \frac{\int_x^1 G(x)\,dx}{\int_0^1 G(x)\,dx} \quad (5)$$

in which

$$G(x) = \exp\left\{-2\int_0^x \frac{M_\delta(y)}{V_\delta(y)}dy\right\} \quad (6)$$

And $v$ is the mutation rate per sequence (or gamete) per generation. As suggested by Kimura [12], the integral with respect to $x$ is over the open interval (0, 1) and actually it is more appropriate if we use $1/(2N_e)$ and $1 - 1/(2N_e)$ as the limit of the integration, especially when the value of the integral changes significantly by including $x = 0$ and 1.

Note

$$-2\int_{x_0}^x \frac{M_\delta(y)}{V_\delta(y)}dy = -4N_e \int_{x_0}^x \left\{-\frac{u_{12}}{1-y} + \frac{u_{21}}{y} + s[h + (1-2h)y]\right\}dy$$

$$= -4N_e \left\{u_{12}\ln(1-y) + u_{21}\ln(y) + s\left[hy + (1-2h)\frac{y^2}{2}\right]\right\}\Big|_{y=x_0}^{y=x}$$

$$= -4N_e \left\{u_{12}\ln\left(\frac{1-x}{1-x_0}\right) + u_{21}ln\left(\frac{x}{x_0}\right)\right.$$

$$\left. + s\left[hx + (1-2h)\frac{x^2}{2} - hx_0 - (1-2h)\frac{{x_0}^2}{2}\right]\right\}$$

Thus,

$$G(x) = \exp\left\{-2\int_{x_0}^{x}\frac{M_\delta(y)}{V_\delta(y)}dy\right\}$$

$$= \exp\left\{-4N_e\left\{u_{12}\ln\left(\frac{1-x}{1-x_0}\right) + u_{21}ln\left(\frac{x}{x_0}\right)\right.\right.$$

$$\left.\left. + s\left[hx + (1-2h)\frac{x^2}{2} - hx_0 - (1-2h)\frac{{x_0}^2}{2}\right]\right\}\right\}$$

$$= \left(\frac{1-x}{1-x_0}\right)^{-4N_eu_{12}}\left(\frac{x}{x_0}\right)^{-4N_eu_{21}}\exp\left\{-4N_es\left[hx + (1-2h)\frac{x^2}{2} - hx_0\right.\right.$$

$$\left.\left. - (1-2h)\frac{{x_0}^2}{2}\right]\right\}$$

$$= C(1-x)^{-4N_eu_{12}}(x)^{-4N_eu_{21}}\exp(-2N_esx[2h + x(1-2h)]) \quad (7)$$

where $x_0 = 1/(2N_e)$ suggested by Kimura and $C$ is a constant,

$$C = (1-x_0)^{4N_eu_{12}}(x_0)^{4N_eu_{21}}\exp\{2N_esx_0[2h + x_0(1-2h)]\} \quad (8)$$

Note $1/[x(1\text{-}x) \times$ Eq. (7)] is the same as Eq. (9.3.4) in [11]. Considering the very low mutation rate per site per generation (i.e., $u_{12}\sim 0$, $u_{21}\sim 0$),

$$C = \exp\{2N_esx_0[2h + x_0(1-2h)]\} \quad (9)$$

$$G(x) = C\exp(-2N_esx[2h + x(1-2h)]) \quad (10)$$

$$\Phi(x) = \frac{2v}{V_\delta(x)G(x)} \frac{\int_x^1 G(x)\,dx}{\int_0^1 G(x)\,dx} = \frac{4N_e v}{x(1-x)G(x)} \frac{\int_x^1 G(x)\,dx}{\int_0^1 G(x)\,dx}$$

$$= \frac{4N_e v}{x(1-x)C\exp(-2N_e sx[2h + x(1-2h)])} \frac{\int_x^1 \exp(-2N_e sx[2h + x(1-2h)])\,dx}{\int_0^1 \exp(-2N_e sx[2h + x(1-2h)])\,dx} \quad (11)$$

Specially, when $h = 0.5$ (note $x_0 = 1/(2N_e)$ suggested by Kimura),

$$C = \exp\{2N_e s x_0\} = \exp(s) \quad (12)$$

$$G(x) = \exp(s)\exp(-2N_e sx) \quad (13)$$

$$\Phi(x) = \frac{2v}{V_\delta(x)G(x)} \frac{\int_x^1 G(x)\,dx}{\int_0^1 G(x)\,dx} = \frac{4N_e v}{x(1-x)\exp(s)\exp(-2N_e sx)} \frac{\int_x^1 \exp(-2N_e sx)\,dx}{\int_0^1 \exp(-2N_e sx)\,dx} \quad (14)$$

Based on Eq. (14), if there is **no selection**, i.e. $s = 0$, we can obtain the well-known site frequency spectrum.

$$\Phi(x) = \frac{4N_e v}{x(1-x)} \frac{1-x}{1} = \frac{\theta}{x} \quad (15)$$

While there is **selection** in Eq. (14)

$$\Phi(x) = \frac{2v}{V_\delta(x)G(x)} \frac{\int_x^1 G(x)\,dx}{\int_0^1 G(x)\,dx} = \frac{4N_e v}{x(1-x)\exp(s)\exp(-2N_e sx)} \frac{\int_x^1 \exp(-2N_e sx)\,dx}{\int_0^1 \exp(-2N_e sx)\,dx}$$

$$= e^{s(2N_e x - 1)} \frac{4N_e v}{x(1-x)} \frac{\exp(-2N_e sx) - \exp(-2N_e s)}{1 - \exp(-2N_e s)}$$

$$= e^{s(2N_e x - 1)} \frac{4N_e v}{x(1-x)} \frac{e^{-2N_e sx} - e^{-2N_e s}}{1 - e^{-2N_e s}} = \frac{4N_e v}{x(1-x)} \frac{e^{-s}\left(1 - e^{-2N_e s(1-x)}\right)}{1 - e^{-2N_e s}} \quad (16)$$

The SFS with or without selection is shown in Fig. S1.

**Note 3. SFS Ratio of Y to Autosome**

To assess the relative selection effect on the Y chromosome compared to Autosome (A), the SFS ratio of Y to A is expressed as F(*x*). Thus, according to Eq. (15) and Eq. (16), the variant frequency of Y chromosome $\Phi'(x)$, assumed to be under selection, is divided to the variant frequency of A, denoted as $\Phi(x)$, which is simply considered as neutral for control:

$$F(x) = \frac{\Phi'(x)}{\Phi(x)} = \frac{N'v'}{Nv} \frac{1}{(1-x)} \frac{e^{-s}\left(1 - e^{-2N's(1-x)}\right)}{1 - e^{-2N's}} \quad (17)$$

in which *x*, *N* and *v* represent the mutation frequency, population size, and mutation rate, respectively. Notably, for Y-linked mutations, these variables are denoted with primes. The relative effective population size of Y-to-A is denoted as $\eta (= N'/N)$, when $\eta < 0.5$, indicating the ratio of $V_Y(K)$ to $V_A(K)$. In addition, the Y-to-A mutation rate, $v'/v$ is equal to 1.68 according to previous research [13, 14]. Then, with substitutions, the Equation (17) is simplified:

$$F(x) = \frac{1.68\eta}{1-x} \frac{e^{-s}\left(1 - e^{-2N\eta s(1-x)}\right)}{1 - e^{-2N\eta s}}$$

Considering the *N* is large and *s* is a negative value of deleterious mutations, and both $e^{-2N\eta s(1-x)}$ $e^{-2N\eta s} \gg 1$ . F(x) in the logarithmic form when x << 1 would be:

$$\ln\left(F(x)\right) \sim \ln\left(\frac{1.68\eta}{1-x} \frac{e^{-2N\eta s(1-x)-s}}{e^{-2N\eta s}}\right)$$

$$= \ln\left(\frac{1.68\eta}{1-x}e^{2N\eta sx-s}\right) \sim [\ln(1.68)+\ln(\eta)-s]+2N\eta sx \quad (18)$$

Therefore, the Equation (18) is a linear function of $x$ with a slope of $2N\eta s$.

**Note 4. The α' value as a function of the distribution of K**

Why would a large $\alpha'$ value necessarily mean "super breeder" males? To see the connection mathematically, it would be simpler to assume $K$ (the progeny number) following a certain mathematical distribution. The gamma distribution has a scale parameter, $\theta$, and a shape parameter, $\kappa$. This flexibility makes it most suited for progeny number distribution.

Given that $K$ follows the gamma distribution, $E_M(K) \sim E_F(K) \sim 1$ over the evolutionary time scale. (Subscript M and F for males and females, respectively). It is also commonly assumed that $K$ in females has $E_F(K) \sim V_F(K) \sim 1$ as in the Wright-Fisher model. Hence, $E_M(K) \sim E_F(K) \sim V_F(K)$. For males, we have

$$E_M(K) = \kappa\theta$$

$$V_M(K) = \kappa\theta^2$$

$$\text{Kurtosis} = 6/\kappa.$$

As defined, $\alpha' = V_M(K)/V_F(K) \sim V_M(K)/E_M(K) = \theta$. Since we observed large $\alpha' > 10$, it means $\theta = \alpha' > 10$. Also, $E_M(K) = \kappa\theta \sim 1$; thus $\kappa < 1/10$. This means $K$ distribution is highly kurtotic with kurtosis > 60. In brief, a large $\alpha'$ value would mean high kurtosis, thus, indicating the $K$ distribution has a very long tail or a very small number of males with very large $K$'s.

Fig. S2 shows the Gamma distribution of $K$ under different $\alpha'$ values (but with the same mean of $E(K) = 1$). It is clear that larger $\alpha'$ values would be associated with a few males having very large breeding successes. Interestingly, the modeling by the Gamma distribution may in fact indicate that the super-breeder males are "outliers" of any mathematical distribution. The main reason is that, under the same distribution, the large $\alpha'$ estimates are also associated with a large proportion of non-breeding males. The existence of outliers in the distribution of breeding successes among males may indicate aspects of reproductive behaviors that are still not understood.

**Note 5. Reduced Y-linked polymorphism: negative vs positive Selection**

This study may also be the first to take into account the two main forces that drive the reported low levels of Y-linked polymorphisms [1, 15, 16], with bonobos displaying the highest diversity on the Y among the great apes [17], consistent with their lowest $\alpha'$. Previous studies have invoked either selection [15, 18, 19] or sex differences in breeding success [16, 20, 21], but not both. Clearly, both forces are operative and, most tellingly, the two forces act multiplicatively in the form of $N_e s$ (see Eq. (6) in methods).

Both positive and negative selection may reduce the level of polymorphism but we focus on negative selection for the following reasons. First, positive selection would have relatively low impact on the low-frequency portion of the spectrum, vis-à-vis negative selection would have [22-25]. More precisely, "the low-frequency portion" does not mean "low-frequency portion" relative to the medium or high-frequency portion of the whole spectrum. It refers to "the relative abundance of low frequency bins from singletons and only up to quadrupletons", which is our primary focus. However, the deviations from the neutral pattern are evident among singleton mutations (see Table 2). Second, negative selection is far more pervasive than positive selection as most mutations are either neutral or deleterious. Third, positive selection reducing polymorphism should be sporadic as sweep via positive selection is transient [26, 27], because the recovery occurs quickly in the low-frequency portion, which is indeed influenced more by negative selection than by positive selection. In contrast, the reduction via negative (background) selection is expected to be nearly constant in time and space. Indeed, Tables 1 and 2 show persistent reductions of Y-linked polymorphism through time.

Finally, the estimation of $\alpha'$ is fundamentally about the nature of genetic drift. JBS Haldane's original model of genetic drift is based on *V(K)* of the branching process [28-30]. Recently, it has been proposed that the WF model should be extended into the WF-Haldane model [31, 32]. In the expanded WF-Haldane model, *V(K)* is the essence of, rather than the add-on to, genetic drift. This study of social-sexual behaviors is a realization of the advantage of the extended WFH model of genetic drift.

### References


1. Hallast P, Maisano Delser P, Batini C *et al.* Great ape Y Chromosome and mitochondrial DNA phylogenies reflect subspecies structure and patterns of mating and dispersal. *Genome Res*. 2016; **26**(4): 427-439. doi: 10.1101/gr.198754.115
2. Nam K, Munch K, Hobolth A *et al.* Extreme selective sweeps independently targeted the X chromosomes of the great apes. *Proceedings of the National Academy of Sciences*. 2015; **112**(20): 6413-6418. doi: 10.1073/pnas.1419306112
3. Xue Y, Prado-Martinez J, Sudmant PH *et al.* Mountain gorilla genomes reveal the impact of long-term population decline and inbreeding. *Science (New York, NY)*. 2015; **348**(6231): 242-245. doi: 10.1126/science.aaa3952
4. Watterson GA. On the number of segregating sites in genetical models without recombination. *Theoretical Population Biology*. 1975; **7**(2): 256-276. doi: 10.1016/0040-5809(75)90020-9
5. Bergström A, McCarthy SA, Hui R *et al.* Insights into human genetic variation and population history from 929 diverse genomes. *Science (New York, NY)*. 2020; **367**(6484). doi: 10.1126/science.aay5012
6. Abecasis GR, Altshuler D, Auton A *et al.* A map of human genome variation from population-scale sequencing. *Nature*. 2010; **467**(7319): 1061-1073. doi: 10.1038/nature09534
7. Delaneau O, Zagury J-F, Robinson MR *et al.* Accurate, scalable and integrative haplotype estimation. *Nature Communications*. 2019; **10**(1): 5436. doi: 10.1038/s41467-019-13225-y
8. Loh PR, Danecek P, Palamara PF *et al.* Reference-based phasing using the Haplotype Reference Consortium panel. *Nature genetics*. 2016; **48**(11): 1443-1448. doi: 10.1038/ng.3679
9. Fu Y-X. Variances and covariances of linear summary statistics of segregating sites. *Theoretical Population Biology*. 2022; **145**: 95-108. doi: https://doi.org/10.1016/j.tpb.2022.03.005
10. Hartl DL, Clark AG, Clark AG. *Principles of population genetics*: Sinauer associates Sunderland, 1997.
11. Crow JF, Kimura M. *An Introduction to Population Genetics Theory*: Blackburn Press, 1970.
12. Kimura M. The Number of Heterozygous Nucleotide Sites Maintained in a Finite Population Due to Steady Flux of Mutations. *Genetics*. 1969; **61**(4): 893-903. doi: 10.1093/genetics/61.4.893
13. Makova KD, Li WH. Strong male-driven evolution of DNA sequences in humans and apes. *Nature*. 2002; **416**(6881): 624-626. doi: 10.1038/416624a
14. Huang W, Chang BH, Gu X *et al.* Sex differences in mutation rate in higher primates estimated from AMG intron sequences. *J Mol Evol*. 1997; **44**(4): 463-465. doi: 10.1007/pl00006166
15. Wilson Sayres MA, Lohmueller KE, Nielsen R. Natural selection reduced diversity on human y chromosomes. *Plos Genet*. 2014; **10**(1): e1004064. doi: 10.1371/journal.pgen.1004064
16. Karmin M, Saag L, Vicente M *et al.* A recent bottleneck of Y chromosome diversity coincides with a global change in culture. *Genome Res*. 2015; **25**(4): 459-466. doi: 10.1101/gr.186684.114
17. Makova KD, Pickett BD, Harris RS *et al.* The Complete Sequence and Comparative Analysis of Ape Sex Chromosomes. 2023: 2023.2011.2030.569198. doi: 10.1101/2023.11.30.569198 %J bioRxiv
18. Rozen S, Marszalek JD, Alagappan RK *et al.* Remarkably little variation in proteins encoded by the Y chromosome's single-copy genes, implying effective purifying selection. *American journal of human genetics*. 2009; **85**(6): 923-928. doi: 10.1016/j.ajhg.2009.11.011
19. Gerrard DT, Filatov DA. Positive and negative selection on mammalian Y chromosomes. *Molecular Biology and Evolution*. 2005; **22**(6): 1423-1432. doi: DOI 10.1093/molbev/msi128
20. Wilder JA, Mobasher Z, Hammer MF. Genetic evidence for unequal effective population sizes of human females and males. *Mol Biol Evol*. 2004; **21**(11): 2047-2057. doi: 10.1093/molbev/msh214
21. Charlesworth B. The effect of life-history and mode of inheritance on neutral genetic variability. *Genet Res*. 2001; **77**(2): 153-166. doi: 10.1017/s0016672301004979

22. Fay JC, Wyckoff GJ, Wu C-I. Testing the neutral theory of molecular evolution with genomic data from Drosophila. *Nature*. 2002; **415**(6875): 1024-1026. doi: 10.1038/4151024a
23. Fay JC, Wyckoff GJ, Wu C-I. Positive and Negative Selection on the Human Genome. *Genetics*. 2001; **158**(3): 1227-1234. doi: 10.1093/genetics/158.3.1227
24. Fay JC, Wu CI. Hitchhiking under positive Darwinian selection. *Genetics*. 2000; **155**(3): 1405-1413. doi: 10.1093/genetics/155.3.1405
25. Wang HY, Chen YX, Tong D *et al.* Is the evolution in tumors Darwinian or non-Darwinian? *National Science Review*. 2018; **5**(1): 15-17. doi: 10.1093/nsr/nwx076
26. Kim Y, Nielsen R. Linkage Disequilibrium as a Signature of Selective Sweeps. *Genetics*. 2004; **167**(3): 1513-1524. doi: 10.1534/genetics.103.025387
27. Kim Y, Stephan W. Joint effects of genetic hitchhiking and background selection on neutral variation. *Genetics*. 2000; **155**(3): 1415-1427. doi: 10.1093/genetics/155.3.1415
28. Haldane JBS. A Mathematical Theory of Natural and Artificial Selection, Part V: Selection and Mutation. *Mathematical Proceedings of the Cambridge Philosophical Society*. 1927; **23**(7): 838-844. doi: 10.1017/S0305004100015644
29. Haldane J. *The causes of evolution*: Longmans, Green, London, 1932.
30. Chen Y, Tong D, Wu CI. A New Formulation of Random Genetic Drift and Its Application to the Evolution of Cell Populations. *Mol Biol Evol*. 2017; **34**(8): 2057-2064. doi: 10.1093/molbev/msx161
31. Ruan Y, Wang X, Hou M *et al.* Resolving Paradoxes in Molecular Evolution: The Integrated WF-Haldane (WFH) Model of Genetic Drift. *bioRxiv*. 2024: 2024.2002.2019.581083. doi: 10.1101/2024.02.19.581083
32. Wang X, Ruan Y, Zhang L *et al.* A Generalized approach to genetic drift and its applications - I. An example of the evolution of ribosomal RNA genes. *bioRxiv*. 2024: 2023.2006.2014.545040. doi: 10.1101/2023.06.14.545040

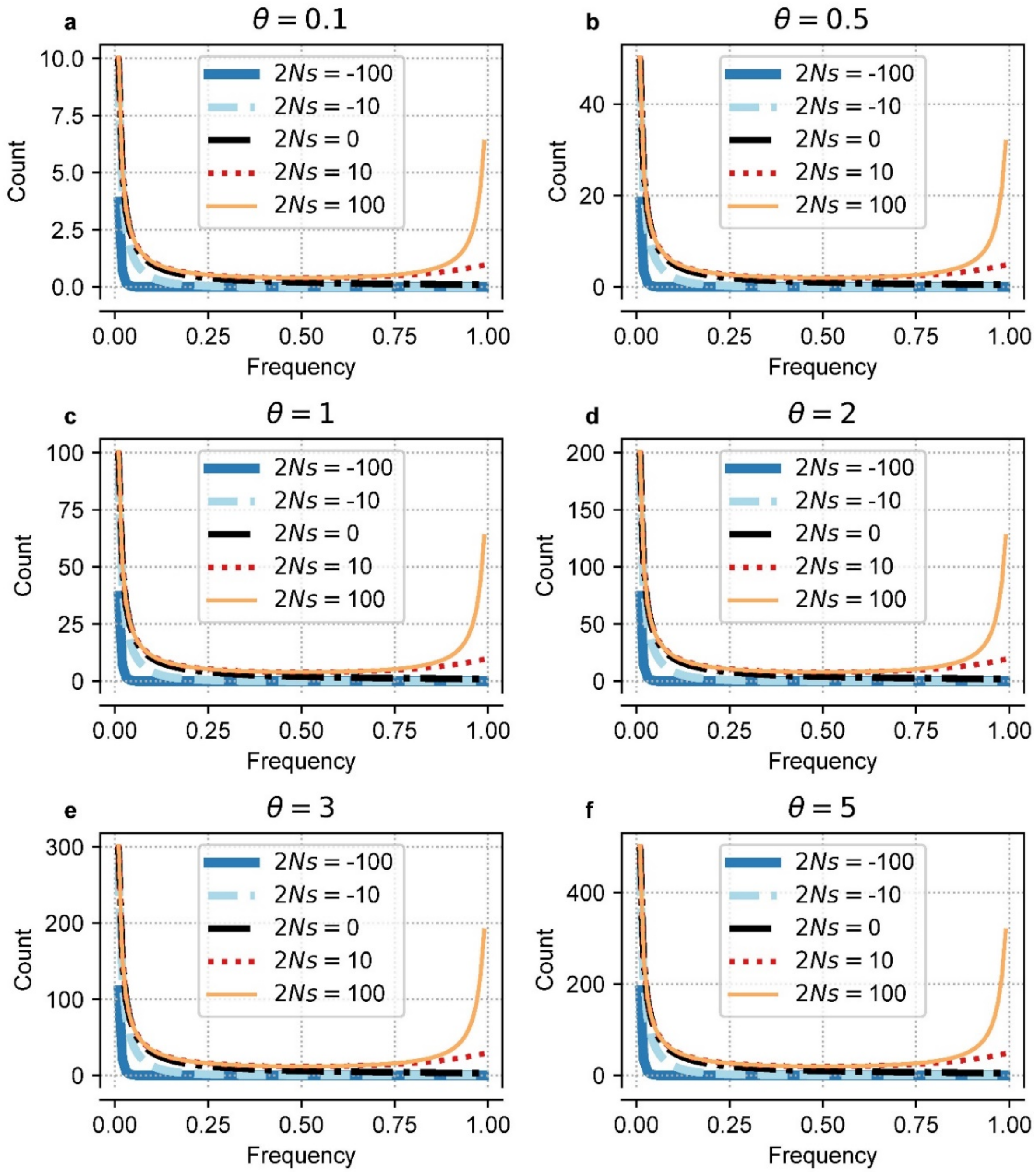


**Fig. S1.** The site frequency spectrum (SFS) with or without selection in a population. The distribution is obtained from an infinite-site model with free recombination, and no dominance. See Eq. (15) and Eq. (16).

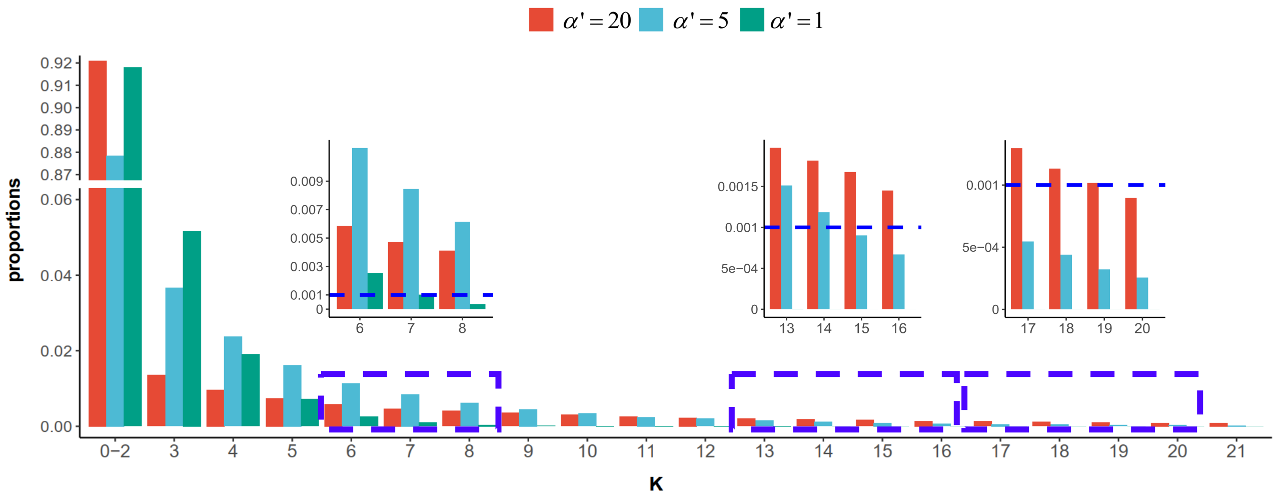


**Fig. S2.** Gamma distribution of $K$ under different $\alpha'$ values. The progeny number, $K$, is governed by three levels of kurtosis determined by $\alpha'$ in the Gamma distribution, with $E(K) = 1$. Three enlarged views display the intervals of $K$ distribution with corresponding blue dashed rectangles. The frequency of $K > 10$ approaches 0 when $\alpha' = 1$, while a large $\alpha'$ value (= 20) is associated with the highest proportion of $K \leq 2$ and $K \geq 20$ among them.

**Table S1.** Summary of genetic variation across different chromosomes in non-human great apes

| | Y Chromosome | | | | X Chromosome | | | | Autosomes | | | |
|---|---|---|---|---|---|---|---|---|---|---|---|---|
| **Common name** | ***N*** | ***S*** | **Total length (bp)** | **$\theta_w$ ($\times 10^{-3}$)** | ***N*** | ***S*** | **Total length (bp)** | **$\theta_w$ ($\times 10^{-3}$)** | ***N*** | ***S*** | **Total length (bp)** | **$\theta_w$ ($\times 10^{-3}$)** |
| **Bonobo** | 4 | 3284 | 3637523 | 0.4924 | 4 | 220 | 268948 | 0.4462 | 26 | 532077 | 160880685 | 0.8667 |
| **Chimpanzee** | | | | | | | | | | | | |
| Western | 9 | 231 | 2496576 | 0.0340 | 7 | 276 | 211364 | 0.5330 | 10 | 738892 | 160880685 | 1.6235 |
| Eastern | 3 | 540 | 2496576 | 0.1442 | 3 | 331 | 211364 | 1.0440 | 12 | 702539 | 160880685 | 1.4460 |
| Nigeria-Cameroon | 4 | 975 | 2496576 | 0.2130 | 4 | 317 | 211364 | 0.8181 | 20 | 401273 | 160880685 | 0.7030 |
| **Gorilla** | | | | | | | | | | | | |
| W lowland | 7 | 689 | 2043299 | 0.1376 | 7 | 128 | 72367 | 0.7219 | 54 | 9983435 | - | 1.5550 |
| E lowland | 3 | 200 | 2043299 | 0.0653 | 3 | 10 | 72367 | 0.0921 | 18 | 3332819 | - | 0.7140 |
| Mountain | 3 | 2 | 2043299 | 0.0007 | 3 | 24 | 72367 | 0.2211 | 14 | 2790350 | - | 0.6520 |
| **Orangutan** | | | | | | | | | | | | |
| Sumatran | 4 | 95 | 2348840 | 0.0221 | 4 | 558 | 247819 | 1.2282 | 10 | 914308 | 160880685 | 2.0089 |
| Bornean | 2 | 318 | 2348840 | 0.1354 | 2 | 138 | 247819 | 0.5569 | 10 | 653958 | 160880685 | 1.4369 |

Note: *N*, number of chromosomes; *S*, number of segregating sites; $\theta_w$, nucleotide diversity based on the number of segregating sites. The Chr X and Chr Y data from [1] ; the autosomes data from [2], except Gorilla autosome data are from [3].

**Table S2.** $\theta$ ($\times 10^{-3}$) for human coding and non-coding regions

Data from HGDP, n = 84 for each population.

| HGDP | $\theta_\pi$ (A) | | | $\theta_\pi$ (X) | | | $\theta_\pi$ (Y) | | |
|---|---|---|---|---|---|---|---|---|---|
| | **Asian** | **European** | **African** | **Asian** | **European** | **African** | **Asian** | **European** | **African** |
| **Coding** | 0.379 | 0.394 | 0.551 | 0.172 | 0.177 | 0.323 | 0.014 | 0.008 | 0.019 |
| **Non-coding** | 0.653 | 0.687 | 0.971 | 0.303 | 0.312 | 0.579 | 0.017 | 0.014 | 0.020 |
| | $\theta_W$ (A) | | | $\theta_W$ (X) | | | $\theta_W$ (Y) | | |
| | **Asian** | **European** | **African** | **Asian** | **European** | **African** | **Asian** | **European** | **African** |
| **Coding** | 0.469 | 0.481 | 0.909 | 0.244 | 0.242 | 0.490 | 0.034 | 0.032 | 0.045 |
| **Non-coding** | 0.719 | 0.738 | 1.430 | 0.383 | 0.384 | 0.850 | 0.052 | 0.036 | 0.053 |

Data from KGP, $n$ = 46 for each population.

| KGP | $\theta_\pi$ (A) | | | $\theta_\pi$ (X) | | | $\theta_\pi$ (Y) | | |
|---|---|---|---|---|---|---|---|---|---|
| | **Asian** | **European** | **African** | **Asian** | **European** | **African** | **Asian** | **European** | **African** |
| **Coding** | 0.373 | 0.396 | 0.524 | 0.146 | 0.169 | 0.265 | 0.012 | 0.005 | 0.002 |
| **Non-coding** | 0.647 | 0.695 | 0.927 | 0.284 | 0.307 | 0.521 | 0.019 | 0.020 | 0.006 |
| | $\theta_W$ (A) | | | $\theta_W$ (X) | | | $\theta_W$ (Y) | | |
| | **Asian** | **European** | **African** | **Asian** | **European** | **African** | **Asian** | **European** | **African** |
| **Coding** | 0.394 | 0.425 | 0.661 | 0.165 | 0.187 | 0.328 | 0.029 | 0.026 | 0.009 |
| **Non-coding** | 0.613 | 0.667 | 1.072 | 0.280 | 0.310 | 0.581 | 0.051 | 0.032 | 0.016 |

Note $\theta_\pi$ is relatively smaller than $\theta_W$ in both datasets, especially in the coding region, implying selection.

**Table S4.** $\alpha'$ estimate based on linear extrapolation for F($x$~0)

| **HGDP (n=84)** | **African** | | | **Asian** | | | **European** | | |
|---|---|---|---|---|---|---|---|---|---|
| | $\Omega$ | **F(0)** | $\alpha'$(YA) | $\Omega$ | **F(0)** | $\alpha'$(YA) | $\Omega$ | **F(0)** | $\alpha'$(YA) |
| $F^*(0)$-F(2/$n$) | 2.7265 | 0.1893 | Inf | 3.6836 | 0.5725 | 0.5793 | 3.1494 | 0.3792 | 1.2414 |
| $F^*(0)$-F(3/$n$) | 3.0682 | 0.2130 | 68.9090 | 1.7784 | 0.2764 | 3.1636 | 2.2970 | 0.2765 | 3.1559 |
| $F^*(0)$-F(4/$n$) | 1.8938 | 0.1315 | Inf | 1.3951 | 0.2168 | 30.8355 | 1.6251 | 0.1956 | Inf |
| F(1/$n$)-F(2/$n$) | 0.6108 | 0.0424 | Inf | 2.5004 | 0.3886 | 1.1759 | 1.4150 | 0.1704 | Inf |
| F(1/$n$)-F(3/$n$) | 1.5406 | 0.1070 | Inf | 1.0180 | 0.1582 | Inf | 1.3149 | 0.1583 | Inf |
| F(1/$n$)-F(4/$n$) | 1.0186 | 0.0707 | Inf | 0.8870 | 0.1379 | Inf | 0.9983 | 0.1202 | Inf |
| F(2/$n$)-F(3/$n$) | 3.8854 | 0.2698 | 3.5120 | 0.4145 | 0.0644 | Inf | 1.2218 | 0.1471 | Inf |
| F(2/$n$)-F(4/$n$) | 1.3154 | 0.0913 | Inf | 0.5283 | 0.0821 | Inf | 0.8385 | 0.1010 | Inf |
| F(3/$n$)-F(4/$n$) | 0.4453 | 0.0309 | Inf | 0.6734 | 0.1047 | Inf | 0.5755 | 0.0693 | Inf |

| **KGP-phase3 (n=46)** | **African** | | | **Asian** | | | **European** | | |
|---|---|---|---|---|---|---|---|---|---|
| | $\Omega$ | **F(0)** | $\alpha'$(YA) | $\Omega$ | **F(0)** | $\alpha'$(YA) | $\Omega$ | **F(0)** | $\alpha'$(YA) |
| $F^*(0)$-F(2/$n$) | 15.0184 | 0.2128 | 74.3664 | 5.2073 | 0.3224 | 1.8678 | 5.4430 | 0.2038 | Inf |
| $F^*(0)$-F(3/$n$) | 4.5076 | 0.0639 | Inf | 2.5864 | 0.1601 | Inf | 4.2620 | 0.1596 | Inf |
| $F^*(0)$-F(4/$n$) | 5.7923 | 0.0821 | Inf | 2.8497 | 0.1765 | Inf | 5.9764 | 0.2238 | 15.2362 |
| F(1/$n$)-F(2/$n$) | 3.8092 | 0.0540 | Inf | 2.0002 | 0.1238 | Inf | 1.3217 | 0.0495 | Inf |
| F(1/$n$)-F(3/$n$) | 1.2437 | 0.0176 | Inf | 1.1297 | 0.0699 | Inf | 1.8585 | 0.0696 | Inf |
| F(1/$n$)-F(4/$n$) | 2.6689 | 0.0378 | Inf | 1.6944 | 0.1049 | Inf | 3.8466 | 0.1440 | Inf |
| F(2/$n$)-F(3/$n$) | 0.4060 | 0.0058 | Inf | 0.6380 | 0.0395 | Inf | 2.6132 | 0.0978 | Inf |
| F(2/$n$)-F(4/$n$) | 2.2340 | 0.0317 | Inf | 1.5595 | 0.0966 | Inf | 6.5622 | 0.2457 | 5.8799 |
| F(3/$n$)-F(4/$n$) | 12.2908 | 0.1742 | Inf | 3.8119 | 0.2360 | 8.0693 | 16.4789 | 0.6170 | 0.5159 |

$\alpha'$ estimations by Y and A chromosomes across three human populations from two datasets reveal the range where the actual $\alpha'$ falls. The $F^*(0)$ used here is set as (1.68×0.5), and the slope between $F^*(0)$ and F(1/$n$) is excluded from the analysis. All the data used here is from Table S3 and summarized in Figure 2.

**Table S3 (separate file).** Low-frequency variants ($\xi_1$ to $\xi_5$) in the site frequency spectrum of each chromosome across two human datasets. Note: $\xi_i$ (= $\theta/i$) is the variant occurring *i* times in each population.

**Table S5 (separate file).** List of human samples from two datasets
